# Late adolescence mortality in mice with brain-specific deletion of the volume-regulated anion channel subunit LRRC8A

**DOI:** 10.1101/2020.05.22.109462

**Authors:** Corinne S. Wilson, Preeti Dohare, Shaina Orbeta, Julia W. Nalwalk, Yunfei Huang, Russell J. Ferland, Rajan Sah, Annalisa Scimemi, Alexander A. Mongin

**Affiliations:** Department of Neuroscience and Experimental Therapeutics, Albany Medical College, Albany, NY, USA; Department of Biomedical Sciences, University of New England College of Osteopathic Medicine, Biddeford, ME, USA; Department of Internal Medicine, Washington University School of Medicine, St. Louis, MO, USA; Department of Biology, University at Albany, State University of New York, Albany, NY, USA

**Keywords:** VRAC, LRRC8A, glutamate-glutamine cycle, GABA signaling

## Abstract

The leucine-rich repeat-containing family 8 member A (LRRC8A) is an essential subunit of the volume-regulated anion channel (VRAC). VRAC is critical for cell volume control, but its broader physiological functions remain under investigation. Recent studies in the field indicate that *Lrrc8a* disruption in brain astrocytes reduces neuronal excitability, impairs synaptic plasticity and memory, and protects against cerebral ischemia. In the present work, we generated the brain-wide conditional LRRC8A knock-out mice (LRRC8A bKO) using *Nestin*^Cre^-driven *Lrrc8a*^flox/flox^ excision in neurons, astrocytes, and oligodendroglia. LRRC8A bKO animals were born close to the expected Mendelian ratio and developed without overt histological abnormalities, but, surprisingly, all died between 5 and 9 weeks of age with a seizure phenotype, which was confirmed by video and EEG recordings. Brain slice electrophysiology detected changes in the excitability of pyramidal cells and modified GABAergic inputs in the hippocampal CA1 region of LRRC8A bKO. LRRC8A-null hippocampi showed increased immunoreactivity of the astrocytic marker GFAP, indicating reactive astrogliosis. We also found decreased whole-brain protein levels of the GABA transporter GAT-1, the glutamate transporter GLT-1, and the astrocytic enzyme glutamine synthetase. Complementary HPLC assays identified reduction in the tissue levels of the glutamate and GABA precursor glutamine. Together, these findings suggest that VRAC provides vital control of brain excitability in mouse adolescence. VRAC deletion leads to a lethal phenotype involving progressive astrogliosis and dysregulation of astrocytic uptake and supply of amino acid neurotransmitters and their precursors.

## 1 INTRODUCTION

The volume regulated anion channel (VRAC) is a chloride/anion channel, ubiquitously expressed in mammalian cells. ^1–3^ The primary accepted physiological role of VRAC is in acute cell volume regulation. Cellular swelling, which occurs due to fluctuations in the extracellular osmolarity or to rises in the concentration of osmotically active molecules within the cell, triggers the opening of VRACs and increases the activity of several types of potassium channels. ^4–6^ The ensuing VRAC-mediated fluxes of Cl^−^ and HCO_3_^−^ provide a countercharge for the loss of intracellular K^+^, leading to the dissipation of osmotic gradients and regulatory cell volume decrease. In addition to inorganic anions, VRAC is permeable to a large variety of small organic osmolytes, which are either negatively charged *(e.g.*, the amino acids glutamate and aspartate, and the carbohydrates, lactate and pyruvate), or net-neutral (*e.g.*, the amino acids alanine and glutamine, and the polyols, sorbitol and myo-inositol).^1;2;7^ The VRAC-mediated loss of organic osmolytes assists in cell volume control but may also play additional roles in cell-to-cell communication and other processes via the release of biologically active molecules.^6;8–10^

Despite more than three decades of extensive biophysical and functional exploration, the molecular identity of VRAC was revealed only a few years ago. In 2014, two laboratories independently discovered that the leucine-rich repeat-containing protein 8A (LRRC8A) is an essential component of VRAC.^11;12^ LRRC8A belongs to a family of five mammalian proteins (LRRC8A-E) and their paralogues in other vertebrate species.^13^ LRRC8 proteins have sequence and structural homology to the channel and gap junction-forming pannexins and invertebrate innexins.^13^ It has been further established that in order to form fully functional VRAC channels, LRRC8A must partner with at least one additional member from the same family (LRRC8B-E).^12^ Consistent with these findings, subsequent structural and mutagenesis studies confirmed that LRRC8 proteins form a hexameric assembly with one central pore and a relatively broad selectivity for anions and small organic osmolytes.^14–18^ Depending on their subunit composition, LRRC8A-containing VRACs exhibit different biophysical properties, including relative permeability to inorganic anions and organic molecules.^19–21^ The existence of functionally distinct VRAC heteromers accounts for the previously reported cell type- and tissue-specific differences in VRAC properties.

Besides its accepted role in cell volume regulation, VRAC contributes to cell proliferation and migration, regulation of cell excitability and trans-epithelial transport, release of signaling molecules, and intercellular communication.^2;5;7^ Until recently, these additional functions were probed using poorly selective pharmacological blockers and correlative observations on changes in VRAC current densities.^7;9;10^ The discovery of the LRRC8 protein family prompted the use of more powerful genetic tools to identify the functional roles of different VRAC subunits. The first attempt to delete LRRC8A aimed at identifying the role of this protein in the immune system through a global deletion of *Lrrc8a*. This resulted in significant embryonic mortality followed by a rapid demise of any live-born animals due to dramatic growth retardation and multiple organ failure.^22^ Therefore, subsequent studies employed conditional, region- and cell type-specific *Lrrc8a* gene deletion. This approach revealed tissue-specific contributions of VRAC to reproductive function (spermatogenesis), regulation of adipocyte and skeletal muscle cell differentiation and size, insulin signaling, glucose homeostasis, innate immunity and endothelial function.^23–29^ Although the central nervous system (CNS) is highly sensitive to cell volume alterations and VRAC activity^9;30–33^, few studies have analyzed the functional significance of LRRC8 proteins in the brain. Our group uncovered a key role for LRRC8A-containing channels in swelling- and agonist-activated release of excitatory neurotransmitters from astrocytes and explored the molecular composition of the endogenous LRRC8 heteromers in brain cells.^21;34^ Another recent *in vivo* study demonstrated that conditional, *Gfap*^Cre^-driven deletion of *Lrrc8a* in astrocytes leads to reduced hippocampal excitability, impaired long-term potentiation (LTP), spatial and contextual learning deficits, and resistance to cerebral ischemia.^35^

In the present work, we produced a conditional, brain-wide deletion of *Lrrc8a*, using the nestin promoter-driven expression of Cre recombinase. This strategy deletes *Lrrc8a* in neurons, astrocytes, and oligodendrocytes and allows for a comprehensive evaluation of the role of VRAC in the brain. The *Nestin*^Cre/+^;*Lrrc8a*^fl/fl^ mice (here referred to as LRRC8A bKO) developed normally but all died at 5 to 9 weeks of age, while showing changes in hippocampal cell excitability and GABAergic transmission, and overt seizure activity. Thus, our study provides important evidence for the crucial role of VRAC in CNS function via regulation of brain excitability during early-to-late adolescence.

## 2 MATERIALS AND METHODS

### 2.1 Ethics statement

All animal procedures used in the present study were approved by the Institutional Animal Care and Use Committee of Albany Medical College (ACUP #18-04002, #21-03001), and strictly conformed to the Guide for the Care and Use of Laboratory Animals as adopted by the U.S. National Institutes of Health.^36^

### 2.2 Key Resources

All key materials and resources are specified in the text.

### 2.3 Animals

Mice were housed in a temperature, humidity and light-controlled facility on a 12 h light/dark cycle and given free access to food and water. Experiments were performed in age-matched controls of both sexes, divided as evenly as possible between males and females. *Lrrc8a*^fl/fl^ (*Swell1*^fl/fl^) mice were generated as previously described.^23^ Brain-specific *Lrrc8a* knockout mice were produced by breeding *Lrrc8a*^fl/fl^ female mice with commercially available *Nestin*^Cre/+^ male mice (B6.Cg-*Tg(Nes-cre)1^Kln^*/J; Jackson Laboratory stock #003771, RRID: IMSR_JAX:003771), both on a C57BL/6J background. *Nestin*^Cre/+^;*Lrrc8a*^fl/+^ heterozygous males were crossed again with *Lrrc8a*^fl/fl^ female mice to produce *Nestin*^Cre/+^;*Lrrc8a*^fl/fl^ knockout mice and littermate controls. Four possible genotypes, *Lrrc8a*^fl^*^/+^* (abbreviated fl/+), *Lrrc8a*^fl/fl^ (fl/fl), *Nestin*^Cre/+^;*Lrrc8a*^fl/+^ (heterozygous deletion, Het), and *Nestin*^Cre/+^;*Lrrc8a*^fl/fl^ (constitutive knockout, KO) were confirmed by PCR analysis. *Lrrc8a* deletion was verified by PCR amplification across the predicted loxP insertion sites surrounding *Lrrc8a e*xon 3. The *Nestin*^Cre^ transgene insertion into mouse chromosome 12 was validated according to the Jackson Laboratory genotyping protocol.

### 2.4 Behavior testing

To characterize the behavioral phenotype of the brain-specific LRRC8A KO mice, we performed several behavioral assays and compared the results in four phenotypes: fl/+, fl/fl, Het, and KO. Only the open field test (OFT) and the elevated plus maze (EPM) were completed due to the early adolescent lethality observed in the KO animals. Additionally, we were able to evaluate the nest-building behavior in mice, which were individually housed for EEG experiments. All behavior tests were performed in mice of both sexes, with planned post-hoc analysis for potential sex differences whenever possible.

Several cohorts of 6 to 8-wk-old animals of all four genotypes were tested at the same time of day (AM) in conditions of dim-light to facilitate animal activity (∼15 lux). Animals were brought in their home cages from their housing room into the behavior testing room and allowed to acclimate for at least 30 minutes prior to the start of any tests. For the OFT, mice were placed, one at a time, into a quadrant operant chamber containing four identical 50×50 cm cubes and recorded for 10 min. The percentage of time the animal spent in the center or the periphery of the OFT, as well as the total distance travelled, were calculated using ANY-maze software (Wood Dale, IL, USA, RRID:SCR_014289). The next day, the same animals performed the EPM test. One at a time, mice were placed in the center of the EPM apparatus with two open arms and two closed arms (32 cm each), standing 39 cm above the ground. Their activity was recorded for 5 min, and ANY-maze software was used to determine the entries into, and the amount of time spent in the open or closed arms, as well as total distance travelled.

In animals that were individually housed for EEG recordings, we serendipitously found striking differences in the nest-building behavior, which were further evaluated. Mice housed in round 24-cm Plexiglas cages were provided with 10 g of fresh paper bedding once a week. 48 hour later cages were observed for the presence or absence of an organized nest, on a yes-or-no basis. The organized nest was defined as consolidated bedding with a depression in the center. The lack of the nest was defined as bedding material scattered throughout the cage.

### 2.5 Spontaneous seizure activity and EEG seizure recordings

Cohorts of 6-8-week-old mice were observed during the light-cycle hours (7:00 AM to 7:00 PM) for up to 6-8 h per day while performing the pre-determined behavioral testing, as described above. During this time, we noticed seizure activity in several animals and videos of the events were recorded whenever possible. Because of the serendipitous nature of these observations, we did not pre-plan or complete 24-h observations of the animals in behavioral cohorts. For the recorded events, seizure semiology was assessed using the Racine scale^37^ for clonic seizures and a modified brainstem seizure scale validated in our prior work.^38^

In a separate cohort of animals, seizure activity was further quantified with electroencephalography (EEG) and continuous video recording. Five-to-six-week-old mice were anesthetized using isoflurane and secured in a stereotaxic frame. Two stainless-steel surgical screws (J.I. Morris, Oxford, MA, USA) were implanted approximately AP −2.0, ML ±2.0 mm from bregma and served as recording electrodes.^39^ A third screw was implanted in the occipital plate to act as a ground. Individual wires from a 3-wire electrode (P1 Technologies, Roanoke, VA, USA) were wrapped around each screw, and the assembly was secured in place using dental cement (Lang Dental Manuf. Co., Inc., Wheeling, IL, USA). Mice recovered for at least 5 days before being moved into cylindrical plexiglass chambers (24 cm diameter × 25 cm height) and attached to an EEG recording system (A-M Systems, Sequim, WA, USA). EEG and video monitoring were performed until knockout animals died or their littermates reached 10 weeks of age. EEG data were analyzed using DataWave Sciworks (A-M Systems, Sequim, WA, USA) and seizures were identified via visual inspection by trained personnel based on the frequency and amplitude of the firing events in each EEG trace.

### 2.6 Primary astrocyte cultures

Primary cultures of mouse astrocytes were prepared from the brain cortices of newborn (P0-P1) animals of both sexes and all four genotypes. Complete litters were utilized, and tail snips of each pup were taken for genotyping. Each pup’s brain tissue was used for an individual cell preparation. Pups were quickly decapitated, and their brains were harvested in ice-cold sterile Dulbecco’s phosphate-buffered saline without calcium and magnesium (ThermoFisher Scientific, cat. #14190144). Brain cortices were dissected out and cleared of meninges. Cortical tissue was then minced using a sterile surgical blade, mechanically homogenized by pipetting through a one-mL pipette tip, and further digested for 5 min at room temperature (22°C) using a TrypLE Express recombinant protease (ThermoFisher Scientific, cat. #12605010). The dissociated cells were sedimented by centrifugation (900 g for 10 min at 4°C). Pellets were resuspended in Earl’s minimal essential medium (MEM) supplemented with 10% heat inactivated horse serum (HIHS) and 50 U/mL penicillin/streptomycin (all components from ThermoFisher Scientific/Invitrogen, Carlsbad, CA, USA). Cells from each pup brain were plated onto an individual T75 flask pre-treated with poly-D-lysine (Sigma-Millipore, cat. #P6407), to obtain an enriched astrocyte population. The cultured astrocytes were grown for at least 10 days and maintained for up to four weeks at 37°C in a humidified atmosphere containing 5% CO_2_/balance air. The purity of cell cultures was routinely confirmed using immunocytochemistry with the astrocyte marker, glial fibrillary acidic protein (GFAP, Millipore-Sigma, cat. #MAB360, RRID: AB_11212597, 1:400).

### 2.7 Processing of tissue samples

For tissue collection, mice were euthanized with a lethal injection of sodium pentobarbital and perfused transcardially with room temperature PBS. This was followed by perfusion with either ice-cold PBS for western blot and amino acid analyses, or 4% paraformaldehyde in PBS for immunohistochemistry.

For western blotting, brain, heart, kidney, lung, and liver were removed and immediately snap-frozen in liquid nitrogen. Prior to assaying, frozen tissues were homogenized using a mechanical homogenizer (PRO200, PRO Scientific, Oxford, CT, USA) in a tissue lysis buffer containing 150 mM NaCl, 50 mM Tris HCl (pH 8), 0.1% Triton X-100 and 5% protease inhibitor cocktail (ThermoFisher Scientific, cat. #78430). Samples were clarified by a brief centrifugation (10,000 g for 5 min at 4°C) and the supernatants were collected and diluted with 2× reducing Laemmeli buffer (Bio-Rad, Hercules, CA, USA, cat. #1610737). For preparation of cell culture lysates, confluent 60-mm Petri dishes were washed from cell culture media with a Basal medium (for composition see below), lysed in 2% SDS plus 8 mM EDTA, and diluted with 4× reducing Laemmeli buffer (Bio-Rad, cat # 1610747).

For immunohistochemistry, brains were removed and post-fixed in 4% paraformaldehyde (Santa Cruz Biotechnology, Dallas, TX, USA, cat. #sc-281692) for 24 h, 15% sucrose solution for 24 h, and 30% sucrose solution for full cryopreservation (all steps at 4°C). Brains were further mounted on a cryostat specimen stage using Tissue-Tek O.C.T. compound (Sakura Finetek, Torrance, CA, USA), frozen to −20°C and sectioned using a CM3050 cryostat (Leica, Wetzlar, Germany, RRID:SCR_016844) into 25-μm thick sections. Sections were collected into cryoprotectant media containing 30% ethylene glycol and 20% glycerol in Tris buffered saline (TBS) and stored at −20°C until morphological or immunohistochemical staining assays were conducted.

### 2.8 Western blot analyses

Protein expression levels were measured in lysates prepared from whole brain, samples of other organs, or from primary astrocyte cultures. Protein samples in reducing Laemmeli buffer were boiled for 5 min (except for VGAT and GAT-1 analyses), loaded onto 10% Mini-PROTEAN TGX gels (Bio-Rad, cat #4561033), and separated according to a standard SDS-PAGE procedure. Separated proteins were electrotransferred onto a polyvinylidene difluoride membrane (PVDF, Bio-Rad, cat. #1620177). Membranes were blocked for 5 min in Tris-buffered saline containing 0.1% Tween-20 (TBS-T) and 5% milk. They were further incubated overnight in the same blocking buffer with one of the following primary antibodies: mouse monoclonal anti-LRRC8A (Santa Cruz, cat. #sc-517113, dilution 1:500), rabbit polyclonal anti-GAD65/67 (Millipore-Sigma, cat. #G5163, RRID: AB_477019, 1:2,000), rabbit polyclonal anti-VGAT (Invitrogen, cat. #Pa5-27569, RRID: AB_2545045, 1:1000), rabbit polyclonal anti-GAT-1 (Abcam, cat. #ab72448, RRID:AB_1268935, 1:500), rabbit polyclonal anti-GLT-1 (Abcam, cat. #ab41621, RRID:AB_941782, 1:10,000), or rabbit polyclonal anti-glutamine synthetase (Millipore-Sigma, cat. #G2781, RRID:AB_259853, 1:20,000). Blots were washed with TBS-T and probed with the species-matching secondary horseradish peroxidase-conjugated antibody: donkey anti-rabbit (GE Healthcare, cat. #NA932, dilution 1:10,000), or sheep anti-mouse IgG (GE Healthcare, cat #NA931, RRID:AB_772210, 1:10,000) in TBS-T containing 5% milk for 2 h at room temperature. After a final wash, immunoreactivity was measured using ECL reagent (GE Healthcare, cat. #RPN2232) and visualized in a Bio-Rad ChemiDoc Imager (RRID:SCR_019684). For loading controls, membranes were either washed overnight in TBS-T or stripped with Restore buffer (ThermoFisher Scientific, cat. #21059), re-probed with the HRP-conjugated primary anti-β-actin antibody for 20 min (Millipore-Sigma, cat. #A-3854, RRID:AB_262011, 1: 50,000), and the β-actin immunoreactivity was visualized as described above. No β-actin immunoreactivity was detected in heart tissue lysates, so these blots were re-probed with rabbit anti-GAPDH antibody (Millipore-Sigma, cat #G9545, RRID: AB_796208, 1:10000) for 1 h at RT. Protein expression levels were semi-quantitively determined by measuring band intensity using ImageJ software^40^ and further normalization to the β-actin or GAPDH signal from the same membrane.

### 2.9 Neuroanatomical analysis and immunohistochemistry

To analyze brain morphology, brain sections from 6-week old animals of all genotypes were stained with thionin. Frozen sections were washed from cryoprotectant media with PBS, mounted on charged slides, and dried overnight. The tissue was progressively rehydrated by dipping into 100%, 95%, 70%, and 50% ethanol, followed by a 15-sec thionin staining (0.25% thionin, 1 M sodium acetate and 1 M glacial acetic acid, pH 4.3). Slides were washed in water and then progressively dehydrated again through the same ethanol solutions before being cleared with Xylene and mounted with Permount (ThermoFisher Scientific, cat. #SP15-100). Thionin-stained brain sections were scanned using a Zeiss AX10 microscope (Carl Zeiss, Oberkochen, DE) at 200× magnification, and the resulting images stitched together with Neurolucida 11.03 software (MBF Bioscience, Williston, VT, USA, RRID:SCR_001775).

For immunohistochemical detection of cell-specific markers, floating sections from the brains of 8-week-old animals were washed from cryoprotectant with PBS and blocked for 1 h at room temperature in PBS with either 3% normal goat serum plus 0.3% Triton X-100 (TX-100) for staining myelin basic protein (MBP), or 2% normal goat serum and 0.2% TX-100 for detecting NeuN, GFAP, parvalbumin, and glutamine synthetase (GS) immunoreactivity. Floating sections were next incubated overnight at 4°C with one or two primary antibodies in blocking buffer with the following dilutions: rabbit polyclonal anti-RBFOX3/NeuN (Novus Biologicals, cat. #NBP1-77686, RRID: AB_11009597, 1:200), mouse monoclonal anti-GFAP (Millipore-Sigma, cat. #MAB360, RRID: AB_11212597, 1:400), mouse monoclonal anti-MBP antibody (Biolegend, cat. #SMI-99P, RRID: AB_10120129, 1:1000), guinea pig polyclonal anti-parvalbumin antibody (Immunostar, cat. #24428, RRID:AB_572259, 1:1000), or rabbit polyclonal anti-glutamine synthetase antibody (Millipore-Sigma, cat. #G2781, RRID:AB_259853, 1:5000). Sections were washed in PBS, and then incubated for 2 h at room temperature with one of the species-matching conjugated secondary antibodies (all from ThermoFisher Scientific): goat anti-mouse Alexa-Fluor 555 (cat. #A-21422, RRID:AB_2535844, 1:400 for MBP staining), donkey anti-mouse Alexa-Fluor 594 (cat #A-32744, RRID:AB_2762826, 1:400 for GFAP or glutamine synthetase), goat anti-guinea pig Alexa-Fluor 488 (cat #A-11073, RRID:AB_2534117, 1:500 for parvalbumin), or goat anti-rabbit Alexa-Fluor 488 (cat # A32731, RRID:AB_2633280, 1:400 for NeuN or GFAP). Sections were washed from secondary antibody with PBS, counterstained with 0.05 μg/mL DAPI at room temperature for 5 minutes, and further washed with PBS. They were mounted on charged slides, dried in the dark overnight at room temperature, and mounted with Vectashield HardSet Antifade Mounting Medium (Vector Laboratories, cat. #H-1400-10).

Immunolabeled brain sections were scanned using a Zeiss AX10 microscope at 200× magnification, and the resulting images stitched together with Neurolucida 11.03 software. Alternatively, images were acquired using a Zeiss LSM880 confocal microscope with a LCI Plan-Neofluar 25x/0.8 Imm Korr DIC objective lens with a 0.761 μm interval through a 5.3 μm z-depth of tissue, stitching together 12 adjacent tiles with 5% overlap, and processed with Zen Software (Carl Zeiss, RRID:SCR_013672). Cortical NeuN-positive cells were quantified in Neurolucida-scanned images, within a predefined ∼2.1 mm^2^ region of cortex situated above hippocampus in sagittal brain sections between +1.7 and +1.9 mm AP from bregma, using ImageJ automatic particle counting.^40^ Tightly packed NeuN-positive cells in the CA1 pyramidal cell layer were quantified in z-stack confocal images using 3D reconstruction and the spot detection function in IMARIS 9.5 software (Bitplane, Zürich, CH, RRID:SCR_007370). For this latter analysis, the spot size was 5.5 and the average analyzed volume was 5.61×10^5^ μm^3^. GFAP-positive cells in the CA1 stratum radiatum layer were quantified in images assembled with Neurolucida software using IMARIS spot detection (spot size of 20, and average analyzed area of 1.31×10^7^ μm^2^). Representative high-magnification confocal images of GFAP-positive cells were acquired using an Alpha Plan-Apochromat 63x/1.46 Oil Korr M27 objective lens with a 0.385 μm interval through an 11.5 μm z-depth of tissue, with adjacent tiles stitched together with 5% overlap.

MBP immunoreactivity was analyzed as the thickness and length of corpus callosum in brain sections from +0.38 to +0.98 mm AP from bregma. We further quantified the MBP signal intensity in several representative ∼0.2 mm^2^ (50 x 50 pixel) regions of the corpus callosum, cerebral cortex, and striatum.

Parvalbumin-positive interneurons were counted in the pyramidal cell layer of the CA1 hippocampal region, or a ∼2.4 mm^2^ region of the cerebral cortex from the midline to a vertical marker at the turn of the hippocampus, in sections between +1.7 and +1.9 mm AP from bregma. Cells were counted manually in the pyramidal cell layer or using automatic particle counting in the cerebral cortex, using ImageJ.

### 2.10 VRAC activity assay

Functional activity of the volume-regulated channel was evaluated by measuring the swelling-activated release of the non-metabolizable analog of glutamate, D-[^3^H]aspartate, as previously described. ^21;41^ Two-to-three-week-old primary astrocyte cultures from all four genotypes were re-plated onto 18×18 mm glass coverslips treated with poly-D-lysine, grown to 80–90% confluency, and loaded overnight with MEM +HIHS cell culture media supplemented with 4 μCi/mL D-[2,3-^3^H]aspartate (Perkin Elmer, Waltham, MA, cat. #NET501001MC). Coverslips were washed in isosmotic Basal medium containing (in mM) 135 NaCl, 3.8 KCl, 1.2 MgSO_4_, 1.3 CaCl_2_, 1.2 KH_2_PO_4_, 10 HEPES, 10 D-glucose (pH 7.4, osmolarity 290 ±2 mOsm) and placed in a custom-made Leucite perfusion chamber with inlet/outlet ports and ∼200 μm of space between the cells and the Teflon screw-top. Swelling was induced by perfusing cells with medium in which osmolarity was decreased to 200 ±2 mOsm (50 mM reduction in [NaCl]). Coverslips were superfused with either isosmotic or hypoosmotic media at a rate of ∼1.2 mL/min. One-minute perfusate fractions were collected into scintillation vials using an automated fraction collector Spectra/Chrom CF-1 (Spectrum Chemical, New Brunswick, NJ, USA). After the last minute of collection, total cell lysates were collected from coverslips using 2% sodium dodecyl sulfate (SDS) plus 8 mM EDTA lysis buffer. Samples were mixed with an Ecoscint A scintillation liquid (Atlanta Biologicals, Atlanta, GA, USA, cat. #LS-273), and [^3^H] content was measured offline in a TriCarb 4910 TR scintillation counter (Perkin Elmer). Results are reported as fractional release rates relative to the total [^3^H] content at each time-point, using an Excel-based custom program.

### 2.11 Electrophysiology

Transverse hippocampal slices (250 μm thick) were prepared from 5-7-week-old C57BL/6J mice of either sex, using a vibrating blade microtome (VT1200S, Leica Microsystems, RRID:SCR_018453). The two genotypes, fl/+ and fl/fl, were used as a combined control and compared to LRRC8A bKO. Mice were deeply anesthetized with isoflurane and transcardially perfused with ice-cold solution containing (in mM): 119 NaCl, 2.5 KCl, 0.5 CaCl_2_, 1.3 MgSO_4_ꞏH_2_O, 4 MgCl_2_, 26.2 NaHCO_3_, 1 NaH_2_PO_4_, and 22 D-glucose (320 mOsm, pH 7.4, pre-equilibrated with 95% O_2_/5% CO_2_). The brain was rapidly removed and placed in ice-cold slicing solution containing (in mM): 227 sucrose, 2.5 KCl, 2 MgSO_4_, 1 CaCl_2_, 1.2 NaH_2_PO_4_, 26 NaHCO_3_, 22 D-glucose (320 mOsm, pH 7.4, constantly bubbled with 95% O_2_/5% CO_2_). Once cut, the slices were stored at 36°C for 30 min and then at room temperature (for up to 5 h), in a submersion chamber containing the same solution used for the transcardial perfusions mentioned above.

Unless otherwise stated, the recording solution contained (in mM): 119 NaCl, 2.5 KCl, 1.2 CaCl_2_, 1 MgCl_2_, 26.2 NaHCO_3_, 1 NaH_2_PO_4_, 22 glucose (pH 7.4). The same recording solution was used to fill the glass capillaries used for extracellular field recordings (R_el_ ∼1.5 mOhm). Stimulating and recording electrodes were placed in the middle third of the hippocampal stratum radiatum, ∼100 μm away from each other. Postsynaptic responses were evoked by delivering constant voltage electrical pulses (50 μs) through a stimulating bipolar stainless-steel electrode (Frederick Haer, Bowdoin, ME, USA, catalog #MX21AES(JD3)). Whole-cell, voltage-clamp patch-clamp recordings were obtained using a pipette solution containing (in mM): 120 CsCH_3_SO_3_, 10 EGTA, 2 MgATP, 0.2 NaGTP, 5 QX-314Br, 20 HEPES (290 mOsm, pH 7.2; R_el_ ∼5-6 MOhm). Current clamp recordings were obtained by replacing CsCH_3_SO_3_ with equimolar KCH_3_SO_3_ (QX-314 was omitted). Data were discarded if the series resistance changed >20% during the course of the experiment. All recordings were obtained using a Multiclamp 700B amplifier and filtered at 10 KHz (Molecular Devices, San Jose, CA, USA, RRID:SCR_018455), converted with an 18-bit 200 kHz A/D board (HEKA, Holliston, MA, USA), digitized at 10 KHz, and analyzed offline with custom-made software (A.S.) written in IgorPro 6.36 (Wavemetrics, Portland, OR, USA, RRID:SCR_000325). All recordings were performed at room temperature.

For multiple-probability fluctuation analysis (MPFA), we recorded inhibitory postsynaptic currents (IPSCs) at different extracellular calcium concentrations (0.5-4 mM). MPFA provides a method for estimating the quantal parameters *N* and *Q* from the IPSC variance and mean, under a range of release probability conditions.^42–44^ The mean IPSC amplitude and variance were normalized to corresponding values measured at [Ca^2+^] =0.5 mM. The resulting data were fit parabolically using the equation 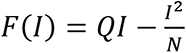.

### 2.12 HPLC analysis of brain amino acid content

Whole-brain tissue lysates were prepared from snap-frozen tissue by homogenizing one hemisphere in 0.4 M perchloric acid. The tissue homogenates were clarified by centrifugation (10,000 g for 10 min). Pellets were saved to determine total protein content and the supernatants were used to determine tissue amino acid levels by reverse-phase high pressure liquid chromatography (HPLC) as previously described.^45^ An Agilent 1200 HPLC setup (RRID:SCR_018037) and Eclipse XDB-C18 column (both from Agilent Technologies, Santa Clara CA, USA) were used for HPLC analysis. Each sample was subjected to a precolumn derivatization with a freshly prepared mix of o-phthalaldehyde and 2-mercaptoethanol in 0.4 M sodium tetraborate buffer (pH 9.5). Derivatized amino acids were eluted from a column with a solvent containing 30 mM NaH_2_PO_4_, 1% tetrahydrofuran, 30 mM sodium acetate, 0.05% sodium azide, and increasing concentrations of HPLC grade methanol (10-30%). A programmable 1200 series fluorescence detector (Agilent) was used to measure the fluorescence signal from each sample. The identification of peaks and concentrations of individual amino acids in the samples were determined by processing amino acid standards of L-alanine, L-aspartate, L-glutamate, taurine, L-glutamine, and GABA. All data were normalized to the total protein content determined in protein pellets after neutralization with 10 N NaOH and sonication in 2% SDS plus 8 mM EDTA using a bicinchoninic acid assay (ThermoFisher Scientific) and bovine serum albumin as a standard.

### 2.13 Statistics

All data are presented as the mean values ±SEM, with the number of independent experiments (n) indicated in each figure. For all experiments, n represents an individual animal as a biological replicate, unless otherwise stated. In radiotracer efflux assays, n represents an individual coverslip (9-12), however all relevant experiments were performed using cells from at least three different cell cultures prepared from different animals.

Experimental group numbers were pre-planned prior to the initiation of experiments, with 10 animals per group for behavioral and HPLC assays, 5 animals per group for western blot analyses, and 6 animals per group for all immunohistochemical and biochemical assays. These group numbers were selected empirically, based on variability seen in our previous studies. Due to the novel nature of this work, specific sample size calculations were done for some but not all experiments. Generally, we planned to be able to detect a difference of 25% between groups with a power of 0.8 and an α of 0.05. In several instances, we performed additional analyses to address reviewers’ questions and the group sizes in western blotting and IHC assays were expanded to 10. In all experiments, animals and samples were coded and analysis was done by a blinded experimenter. A small number of experimental values were excluded from analysis but only for the following reasons: a data point in OFT behavioral assay (fl/fl behavioral time point) was more than 2 SD away from the mean, and a few aberrant values in VRAC activity assays were excluded due to broken coverslips or disrupted perfusion. All data have been included in the primary data files for each figure, and any exclusions from analysis have been noted.

Experimental data were tested for normal distribution. In most cases, statistical differences between groups with normal distribution were determined by one-way ANOVA with Bonferroni post hoc correction for multiple comparisons. If not normally distributed, results were analyzed with the Kruskal-Wallis test and Dunn’s post hoc correction for multiple comparisons. For western blot analyses, optical densities were normalized to β-actin and then again to fl/+ controls in each blot. In this latter instance, we used a Student’s one-sample t-test, compared the mean value in each group to 1, and applied Bonferroni correction for multiple comparisons. Corrected significance values were calculated according to the formula: p_adjusted_ = 1 – (1-p)^n^, where p is the unadjusted p-value and n is the number of comparisons. Fluorescence intensity quantification values (measured in GFAP, MBP, and glutamine synthetase staining) were normalized to the average fluorescence intensity for each immunohistochemistry set of two animals per genotype. These data were analyzed using two-way ANOVA with a fixed effect of genotype and a random effect of set number to remove between-set variability. In all cases, there was no significant effect of set number or interaction between set number and the effect of genotype, therefore all values were graphed together. Electrophysiological studies comparing control and KO slices were analyzed using an un-paired t-test or the non-parametric Mann Whitney test, as appropriate. In EEG recording cohorts, nest-building and seizure phenotypes were scored on a binary basis and further analyzed with the Freeman-Halton extension of Fisher Exact Probability test, using 2×4 contingency tables (www.vassarstats.net). For all figures, *p <0.05, **p <0.01, ***p <0.001, or are not significant unless otherwise noted. All statistical analyses and figures were prepared using Prism 7.0 (GraphPad, San Diego, CA, USA, RRID:SCR_002798).

## 3 RESULTS

### 3.1 Brain-specific deletion of LRRC8A causes 100% mortality in late adolescence

We generated a whole-brain knockout of the essential VRAC protein LRRC8A by utilizing *Nestin*^Cre^-driven deletion of *Lrrc8a*^fl/fl^. Nestin is an intermediate filament protein, which is expressed in developing mouse embryos as early as E7.5 and enriched in neural stem cells throughout development and into adulthood.^46;47^ Nestin promoter-driven Cre targets all classes of brain neuroectodermal cells, including neurons, astrocytes and oligodendrocytes, and is used for brain-specific modifications of gene expression ^48;49^. The breeding scheme and the four genotypes obtained from this strategy are depicted in Fig. 1A. This breeding strategy has been designed to yield a high number of knockout mice and does not generate *Lrrc8a*^+/+^ or *Nestin*^Cre/+^ progeny. We compared the age-matched fl/+, fl/fl, Het, and KO littermates to establish the effect of deletion and reveal potential effects of the fl/fl allele and haploinsufficiency.

**FIGURE 1.**
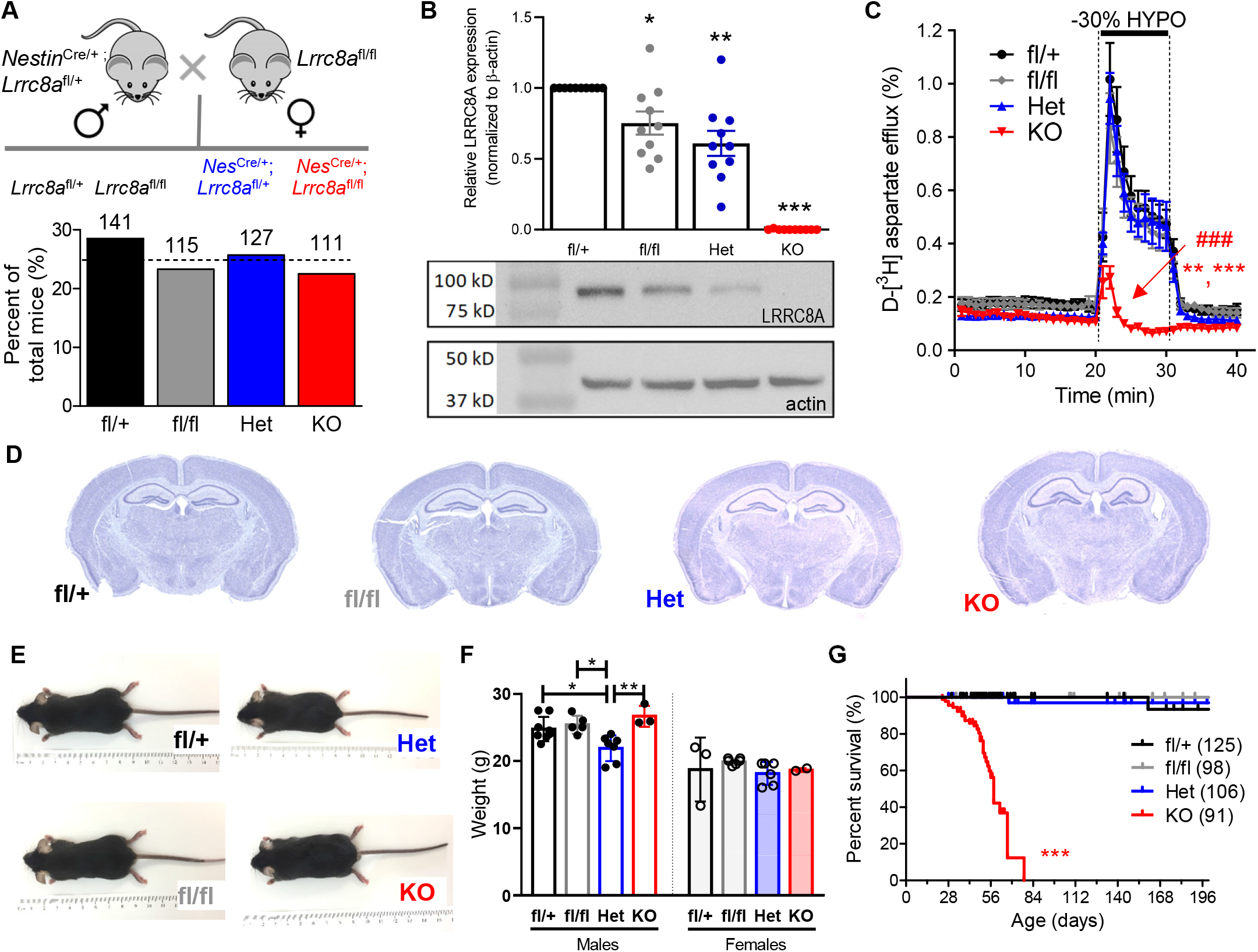
Initial characterization and mortality of animals with the brain-specific deletion of the LRRC8A protein. (A) Breeding strategy and the genotype distribution for the tissue-specific deletion of LRRC8A in the brain using *Nestin*^Cre^-targeted excision of *Lrrc8a*^fl/fl^. The four potential genotypes from this breeding strategy are *Lrrc8a*^fl/+^ (fl/+), *Lrrc8a*^fl/fl^ (fl/fl), *Nestin*^Cre/+^;*Lrrc8a*^fl/+^ (Het) and *Nestin*^Cre/+^;*Lrrc8a*^fl/fl^ (KO) mice. Dotted line denotes expected Mendelian ratio; numbers indicate the total number of animals of each genotype. No significant difference between groups in Chi Square analysis. (B) Western blot analysis of LRRC8A protein levels in whole-brain lysates, normalized to β-actin of each sample and then again to fl/+ controls for each set. Data are the mean values ±SEM from 10 different animals per group. ***p <0.001, one-sample t-test with Bonferroni correction. Lower inset shows representative western blot images for LRRC8A and β-actin re-probed on the same membranes. (C) VRAC activity in primary astrocyte cultures, measured as swelling-activated D-[^3^H]aspartate release in response to hypoosmotic medium. Data are the mean values ±SEM of 9 independent experiments per genotype in three different astrocyte cultures. **p < 0.01, maximum release, KO vs. fl/fl, ***p < 0.001, KO vs fl/+ and Het. ^###^p < 0.001, integral (10-min) release values KO vs. all other groups. ANOVA with Bonferroni correction. (D) Representative thionin-stained brain slices used for neuroanatomical analysis (n =6 per genotype). (E) Representative 8-week old male mice of each genotype. (F) Body weight in male and female animals of all genotypes at 8 weeks. *p<0.05, Het vs fl/+ and fl/fl, **p<0.01, Het vs. KO, two-way ANOVA with Bonferroni correction. Significant effect of sex (***p < 0.001) and genotype (**p < 0.01), no interaction. (G) Kaplan-Meier survival curve of animals of both sexes with the brain deletion of LRRC8A and their littermates. The curve is censored for animal euthanized for tissue analyses. ***p < 0.001, Log-rank (Mantel-Cox) test.

To validate the successful deletion of the targeted protein, we performed semi-quantitative western blot analysis in brain lysates and found a complete absence of LRRC8A immunoreactivity in bLRRC8A KO brains (Fig. 1B; the complete set of full-length western blot images is in Suppl. Fig. 1.1). We also observed moderate reduction in brain LRRC8A protein levels in fl/fl and Het mice (∼25% and ∼40% decreases, respectively, Fig. 1B). As an additional control, we analyzed LRRC8A expression in the heart, kidney, liver, and lung tissues, and found no significant changes in relative LRRC8A protein levels of fl/+ or KO mice (Suppl. Fig. 1.2). We further compared the levels of LRRC8A protein expression in primary astrocyte cultures prepared from neonatal brains of all four genotypes and confirmed the loss of LRRC8A in the LRRC8A KO cells (Suppl. Fig. 1.3). Functional deletion of LRRC8A-containing VRAC was validated in a radiotracer efflux assay, which measures the swelling-activated release of the glutamate analogue D-[^3^H]aspartate from cultured astrocytes exposed to a 30% reduction in medium osmolarity (Fig. 1C). Our prior RNAi experiments in rat astrocytes proved that such release is mediated by the LRRC8A-containing VRAC and represents a functional equivalent of swelling activated Cl^−^ currents ^21;34;41^. The advantage of this approach is that it allows for analysis of VRAC activity in intact cells and on a population rather than a single-cell level. Swelling-activated radiotracer release was significantly reduced in LRRC8A KO astrocytes (∼70% inhibition of the maximal release rates and >80% drop in stimulated integral release values), validating the loss of VRAC activity in LRRC8A KO cells (Fig. 1C). The small residual swelling-activated glutamate release showed different kinetic properties (rapid inactivation, Fig. 1C) and was likely mediated by an alternative, swelling-activated release pathway. Together, these results indicate that there is a brain-specific loss of LRRC8A protein and VRAC activity in bLRRC8A KO mice.

The previously characterized global LRRC8A KO mice show pronounced embryonic lethality, dramatic growth retardation, and die from multiple organ failure.^22^ In contrast, mice with the brain-specific loss of *Lrrc8a* were born close to the expected Mendelian ratio of 25% (22.5%, 111/494 mice, no different, Chi-squared test, Fig. 1A) and developed normally alongside their control (fl/+ and fl/fl) and heterozygous (Het) littermates. We found no gross neuroanatomical or cytoarchitectural abnormalities in the forebrain of 6-week-old animals based on typical cortical layer organization and neuronal morphology in the cerebral cortex and hippocampus (thionin staining in Fig. 1D, see Suppl. Fig. 1.4 for extended analysis). We did not detect any gross defects in body anatomy (Fig. 1E) or reductions in body weight in KO mice of either sex at 8-weeks of age (Fig. 1F).

Despite the lack of overt abnormalities in brain structure or body size, group-housed LRRC8A bKO mice began dying between postnatal weeks 4 and 5 and all remaining knockouts died by the end of postnatal week 9, which approximately corresponds to puberty and late adolescence, respectively^50^ (see the Kaplan-Meier survival curve in Fig. 1G, note that the curve is censored for animals euthanized for experimental tissue collection at 6 and 8 weeks of age). Gross anatomical evaluation did not reveal any obvious structural abnormalities, bleeding in the brain or defects of major organs in the chest cavity that could account for a cause of death for bLRRC8A KO mice. The possible reasons for mortality are explored below.

### 3.2 LRRC8A bKO mice display astrogliosis, but no changes in numbers of neurons and astrocytes, and myelination

As the *Nestin*^Cre^-driven deletion of *Lrrc8a* targets all cells derived from neural precursors, we examined whether VRAC deletion leads to changes in the numbers of neurons, astrocytes, and/or oligodendrocytes. To this end, we used immunohistochemical analysis and counted the numbers of cells immunolabelled for the neuronal marker NeuN (Fig. 2A), the astrocytic marker GFAP (Fig. 2B), and the oligodendroglial marker MPB (Fig. 2C) in 8-week-old brains. In all cases, the statistical comparisons were done across all four genotypes.

**FIGURE 2.**
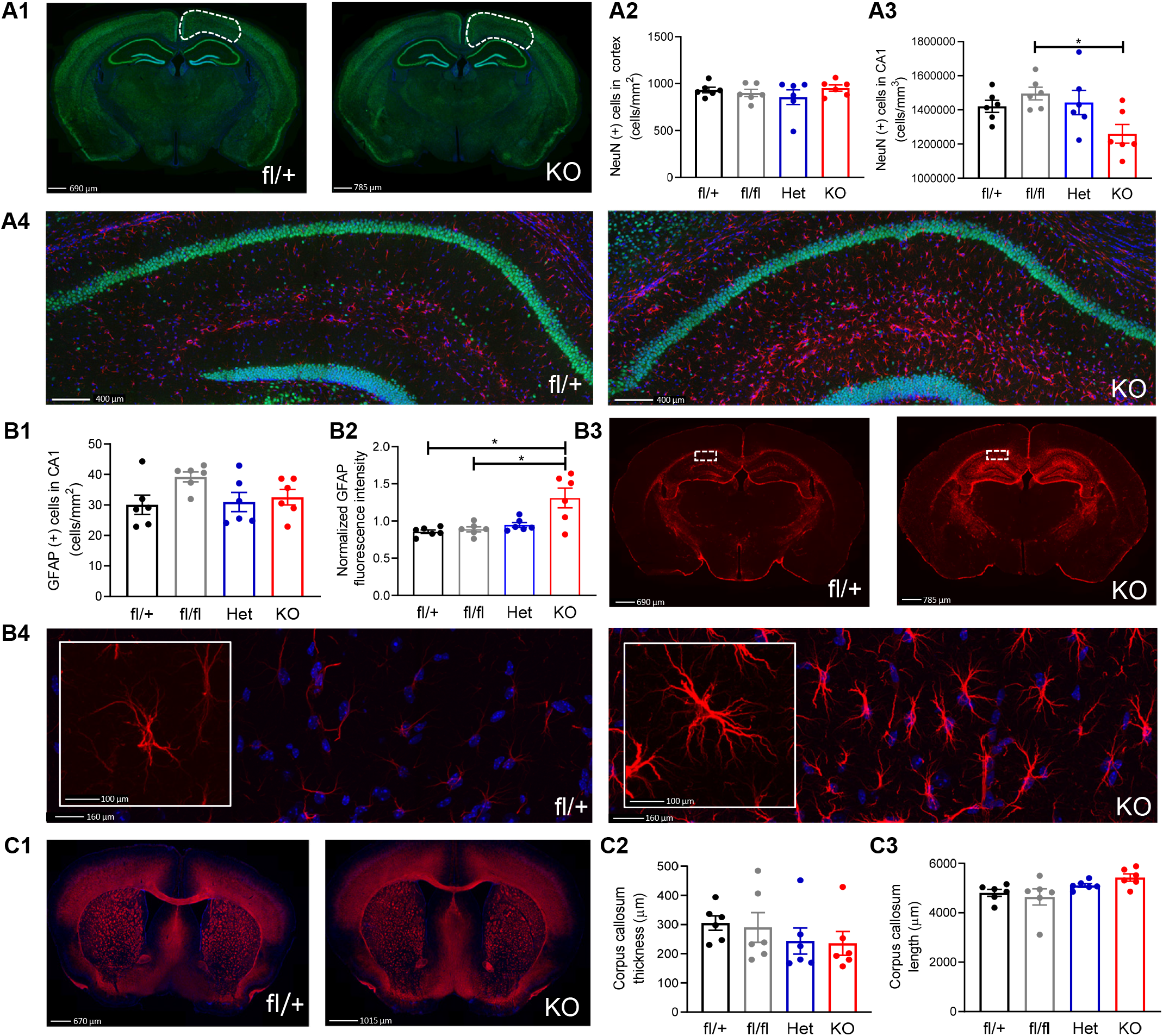
Immunohistochemical analysis of brain cell populations targeted by the *Nestin*^Cre^ promoter. (A1) Representative images of the immunoreactivity for neuronal marker NeuN (green) and DAPI (blue) in 8-week old fl/+ controls (left) and KO (right) brains. (A2) Quantification of NeuN+ cell numbers in the cortical region outlined in A1. (A3) Quantification of NeuN+ cell numbers in the CA1 pyramidal cell layer, shown in A4. Data are the mean values ±SEM from 6 animals per group, analyzed by one-way ANOVA with Bonferroni correction. *p < 0.05, fl/fl vs. KO. (A4) Representative high-magnification images of the immunoreactivity for NeuN (green), the astroglial cell marker GFAP (red), and DAPI (blue) in fl/+ controls (left) and KO (right) brains. (B) Quantification of GFAP+ cell numbers (B1) and the relative GFAP immunofluorescence normalized to the average intensity of each set (B2) in hippocampi of four genotypes. Data are the mean values ±SEM of 6 animals per group, analyzed by one-way ANOVA (B1) or two-way ANOVA (B2) with Bonferroni correction. No significant effect of set number. *p < 0.05, effect of genotype, fl/+ and fl/fl intensity vs. KO. (B3) Representative images of GFAP (red) in hippocampal areas of fl/+ controls (left) and KO (right) animals. (B4) Representative high-magnification images of GFAP (red) and DAPI (blue) immunoreactivity in the CA1 stratum radiatum region outlined in B3. High magnification insets more clearly show astrocyte morphology in each of these genotypes. (C1) Representative immunoreactivity and quantification of the oligodendroglial marker, myelin basic protein (MBP, red) and DAPI (blue) in fl/+ controls (left) and KO (right) brains. (C2-C3) Quantification of corpus callosum medial thickness (C2) and average length (C3) in four genotypes. Data are the mean values ±SEM of 6 animals per group, analyzed by Kruskal-Wallis with Dunn’s test (C2) or one-way ANOVA with Bonferroni correction (C3).

The number of NeuN-positive cells was quantified in a defined, representative region of interest in the cerebral cortex above the hippocampal CA1 region and showed no differences across the four genotypes (Fig. 2A2). Analysis of hippocampal CA1 pyramidal cell layer in the same brains showed a slight decrease in NeuN-positive cells in bLRRC8A KO brains (11-15% lower than in three other genotypes, significant vs. fl/fl only).

There were no changes in the numbers of GFAP-positive astrocytes, quantified in the CA1 region of the hippocampus (Fig. 2B1). Due to low GFAP expression in the cerebral cortex, counts of cortical astrocytes were not performed. Interestingly, although the numbers of GFAP-positive astrocytes in the hippocampus did not differ among the four genotypes, we detected a significant increase in GFAP intensity in bLRRC8A KO specimens, which is often a measure of reactive gliosis (Fig. 2B2, B3 and 2B4). When GFAP immunoreactivity was quantitatively compared with the identical settings and within the sets immunostained at the same time, average normalized intensity levels in bLRRC8A KO sections were increased by >50% (Fig. 2B4), and analysis of LRRC8A bKO specimens at higher magnification revealed hypertrophied astrocytes (inset in Fig. 2B4). Whether this is inherent to bLRRC8A KO or a result of seizure activity is not clear.

To assess oligodendroglia, we stained brain sections for the marker of myelinating cells, MBP, and quantified the thickness and length of the corpus callosum, a major myelinated tract in the brain (Fig. 2C). We found no changes in average corpus callosum thickness (Fig. 2C2) or length (Fig. 2C3) in bLRRC8A KO mice. The average fluorescence intensity of MBP in the cerebral cortex or striatum did not differ (data not shown, available in the Open Science Framework).

Overall, these findings suggest that deletion of *Lrrc8a* does not markedly affect the numbers of neurons and astrocytes, or the degree of myelination. The lack of changes in the CNS was somewhat surprising because VRAC has been proposed to regulate cell proliferation, migration, differentiation, and survival in many cell types (reviews^2;4;5^ and recent experimental studies^22;25;51^).

### 3.3 Behavioral changes in LRRC8A bKO mice

We preplanned several behavioral assays on mice of both sexes, aged 6-7 weeks, but only two assays were completed due to the early onset of animal mortality. The final resulting distribution among sexes was as follows: fl/+: 6 males, 3 females; fl/fl : 3 males, 7 females; Het: 5 males, 5 females; and KO: 5 males, 5 females. We used the open field and elevated plus maze tests to measure locomotion and anxiety-like behaviors in control and KO mice (see Fig. 3 and extended analysis in Suppl. Fig. 3.1). In the open field test, LRRC8A bKO animals spent significantly less time in the center of the open field arena (Fig. 3A, B), but displayed no reduction in locomotor activity (Fig. 3C), suggesting an anxiety phenotype. However, this conclusion was not supported by the outcomes of the elevated plus maze test. In the latter assay, LRRC8A bKO mice did not show any significant decrease in the number of entries into the open arms and thus did not display an anxiety-like phenotype as compared to controls and the animals with heterozygous LRRC8A deletion (Fig. 3D, E). Again, there were no changes in the total distance travelled between all four genotypes (Fig. 3F). Of note, fl/fl animals also spent significantly less time in the center of the open field test compared with control fl/+ mice, however it is unclear whether this demonstrates a real phenotype.

**FIGURE 3.**
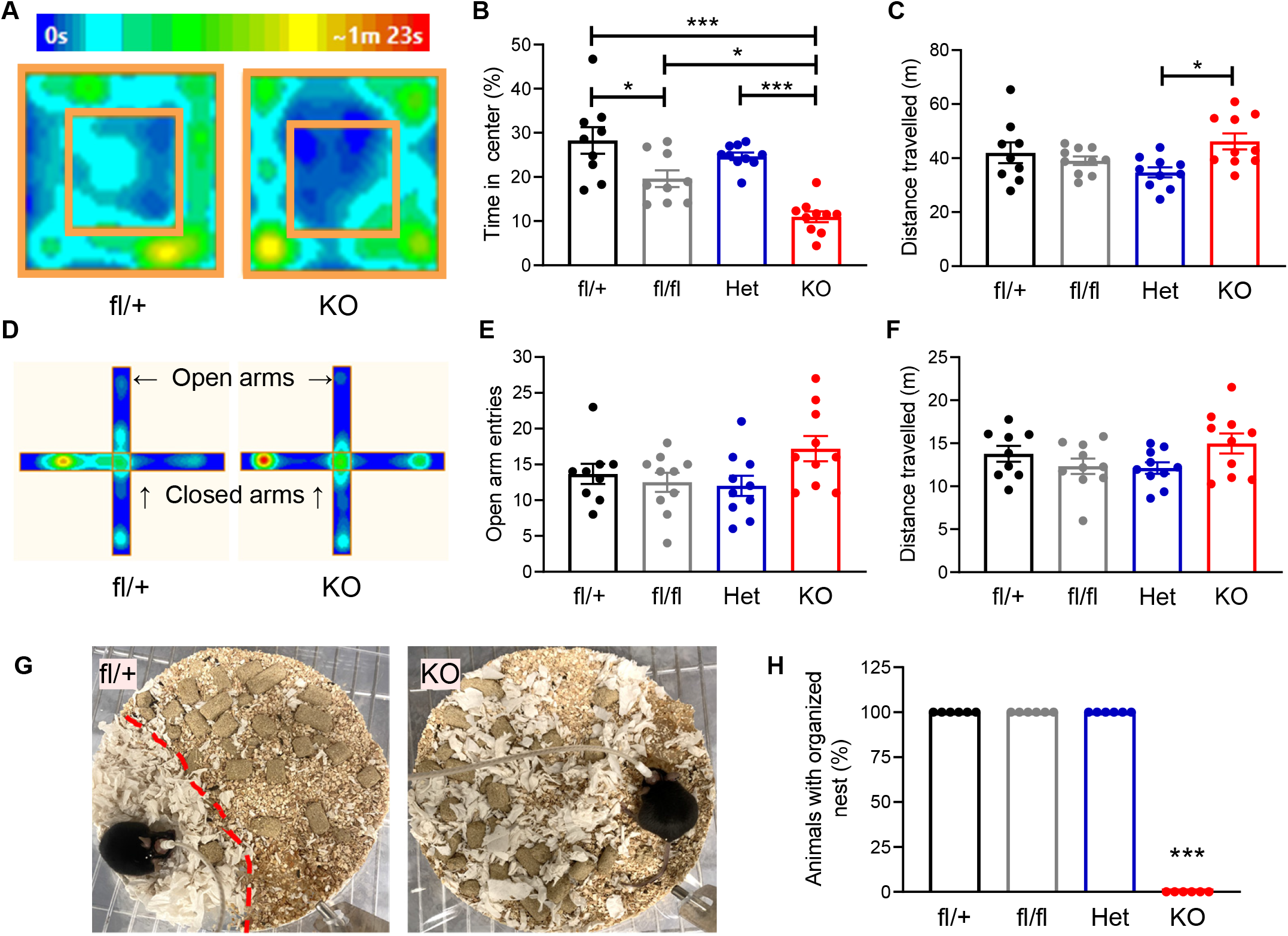
Impact of brain-specific LRRC8A deletion on animal behavior. (A) Representative heat map for fl/+ control and bLRRC8A KO mice in the open field test. Color heat map indicates the amount of time spent in the center or periphery of the operant chamber. (B) Percentage of the total time that the animals of four genotypes spent in the center of the open field. Data are means ±SEM of 9 fl/+ and fl/fl mice and 10 mice in other groups, in mice of both sexes. *p <0.05, fl/+ vs. fl/fl and fl/fl vs. KO; ***p < 0.001, fl/+ vs. KO and Het vs. KO, analyzed by one-way ANOVA with Bonferroni correction. (C) Total distance traveled during the open field test. Data are means ±SEM of 9 fl/+ mice and 10 mice in other groups. *p < 0.05, Het vs. KO, one-way ANOVA with Bonferroni correction. (D) Representative heat maps for fl/+ controls and KO mice in the elevated plus maze with two open and two closed arms. (E) Number of entries into the open arm of the elevated plus maze. Data are the means ±SEM of 9 fl/+ mice and 10 mice in other groups. (F) Distance traveled during the elevated plus maze trial in the same animals as in E. (G) Representative images of nest building and maintenance behavior of fl/+ (left) and KO (right) mice captured 48 h after providing fresh nesting material. (H) Summary of nesting phenotypes in the four animal cohorts used in the EEG experiments. Presence of the nest was scored on a binary basis (yes-or-no). ***p < 0.001, KO vs. all other genotypes, Fisher Exact Probability test with the Freeman-Halton extension.

In the follow-up EEG study, which is described below, separate cohorts of mice (n=6/genotype) were individually housed in plexiglass cages. Serendipitously, we noticed that bLRRC8A KO animals failed to build and maintain a consolidated nest. To explore if this was a genotype-linked phenomenon, we systematically evaluated the presence or absence of an organized nest. Strikingly, we identified a yes-or-no pattern of nest-building, where 100% of bLRRC8A KO mice failed to build a nest (Fig. 3G, H). This behavior was so distinctive that bLRRC8A KO genotype could be visually identified based on nest absence alone.

Overall, behavioral experiments show that deletion of LRRC8A in the brain leads to behavioral changes. There were no differences between male and female mice (post hoc analysis in Suppl. Fig. 3.1).

### 3.4 Seizure activity as a potential cause of animal mortality in bLRRC8A KO mice

During the course of behavioral testing, we noted seizure-like events in bLRRC8A KO animals that were monitored for 4-8 hours a day (Fig. 4A). With the caveat that we did not perform 24-h monitoring, we found that 40% of bLRRC8A KO animals (6/15) displayed an overt seizure phenotype (Fig. 4A). The following two major types of seizures were seen in KO mice: (1) generalized clonic seizures, consisting of forelimb and/or hindlimb clonus with or without loss of posture control, and (2) generalized clonic seizures with loss of posture that immediately transitioned into brainstem-tonic seizures consisting of wild-running and “popcorning” (Suppl. Media File 1, additional video available on the Open Science Framework). No behavioral seizures were ever detected in fl/+, fl/fl, or Het littermates (Fig. 4A).

**FIGURE 4.**
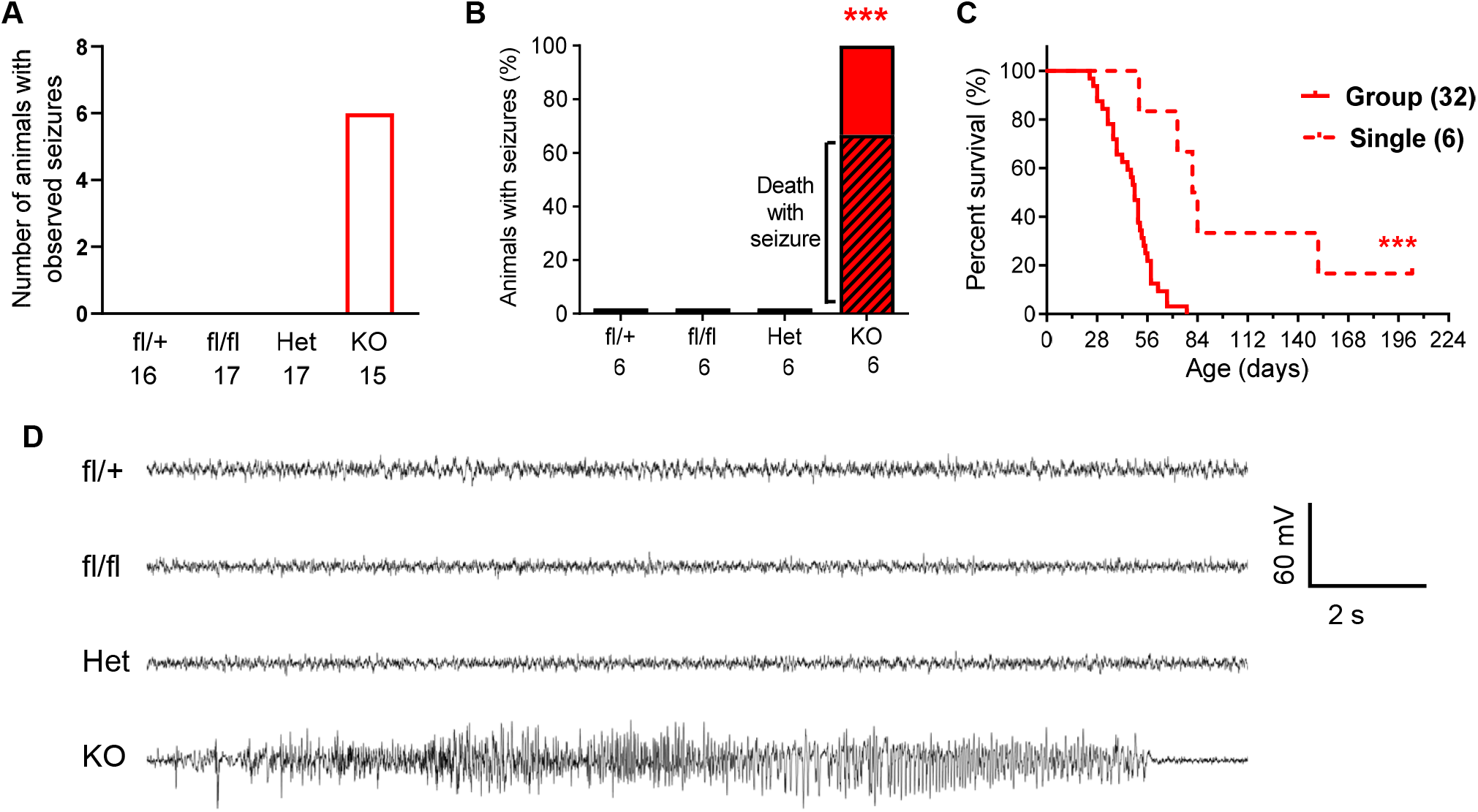
Behavioral and electroencephalographic (EEG) seizures in LRRC8A bKO mice. (A) Number of mice with seizures observed during behavior testing in all four genotypes. Values below each bar represent the total number of animals per group. (B) Percentage of transgenic mice that manifested EEG and behavioral seizure activity during 24/7 monitoring (n=6/group). ***p<0.001, KO vs. all other groups, Fisher Exact Probability test with the Freeman-Halton extension. (C) Comparison of life expectancy in group-housed (n=32) and single-housed (n=6) bLRRC8A KO mice. ***p < 0.01, Log-rank (Mantel-Cox) test, group-housed vs. single-housed. (D) Representative contemporaneous EEG traces from fl/+ (C1), fl/fl (C2), Het (C3), or KO (C4) mice.

The fact that we did not observe seizures in 60% of KO mice or in any of 50 mice in “control” groups (Fig. 4A) does not necessarily mean that these were seizure-free, since the seizure activity could have happened outside of the observation period or was not evident. For this reason, we initiated a separate set of experiments with continuous cortical EEG and video recordings in freely moving animals. Due to time and equipment constraints, the group sizes were limited to 6 animals per genotype (mixed sexes). Two mice were excluded from the analysis due to surgical and post-surgical complications (one fl/+ animal died within 24-h after attachment to the EEG setup, and one moribund KO animal was euthanized). We were able to register multiple characteristic seizure events in 100% of KO mice, which were confirmed by both EEG and video recordings (Fig. 4B, representative EEG traces are shown in Fig. 4D). No seizures were ever detected in the other three genotypes (18 animals and ∼400 recorded days in total, Fig. 4B). Importantly, four out of six KO animals (67%) died with an EEG seizure preceding the moment of death (Fig. 4B). One animal died with a history of seizures but no abnormal EEG activity at the time of death. Finally, one female KO mouse had persistent and numerous seizures but remained alive until the end of the experiment; it was euthanized at the age of 6.8 months (Fig. 4C). The observation that individual housing for EEG recordings (done to avoid animal injury and equipment damage) dramatically extended the lifespan of bLRRC8A KO mice (Figs. 1G and 4C) was unexpected. The significance of this finding and the underlying mechanisms remain unclear. However, it is important to stress that extended life expectancy was not due to experimental or genetic variations because the contemporary group-housed bLRRC8A KO mice continued to die at the initially reported rate.

### 3.5 Electrophysiological analysis in hippocampal slices reveals reduced excitability of CA1 pyramidal cells in LRRC8A bKO mice

Since spontaneous seizure activity was observed in mice with brain-specific deletion of *Lrrc8a*, we asked whether this could be due to changes in cell excitability. We looked at neuronal excitability in hippocampus, a brain region implicated in seizure generation. ^52;53^ To this end, we performed whole-cell patch-clamp recordings from CA1 pyramidal cells (CA1-PCs) in acute hippocampal slices from 5-7 week-old KO mice. Fl/+ and fl/fl mice were used as combined controls. Our first experiments aimed to identify changes in the passive and active membrane properties of CA1-PCs (Fig. 5A,B), the main output neurons of the hippocampus. There were no statistically significant changes in the membrane resistance (Fig. 5B1) or resting membrane potential across genotypes (Fig. 5B2). The holding current necessary to keep these cells at a potential of −70 mV was similar across all tested genotypes (Fig. 5B3). In separate current clamp experiments, we maintained all cells at −70 mV using DC current injections and delivered 5-ms positive current steps of increasing amplitude to evoke single action potentials to measure the rheobase (Fig. 5C, D). The rheobase, which represents the amplitude of the smallest current step able to elicit an action potential, was larger in CA1-PCs from LRRC8A bKO mice (Fig. 5D1). The threshold for action potential generation was more depolarized in KO group (Fig. 5D2), as was the peak of the action potential (Fig. 5D3). The action potential duration, measured from the full width at half maximum (FWHM), was not changed (Fig. 5F1). These findings suggest the existence of a subtle decrease in the excitability of CA1-PCs in the absence of LRRC8A protein.

**FIGURE 5.**
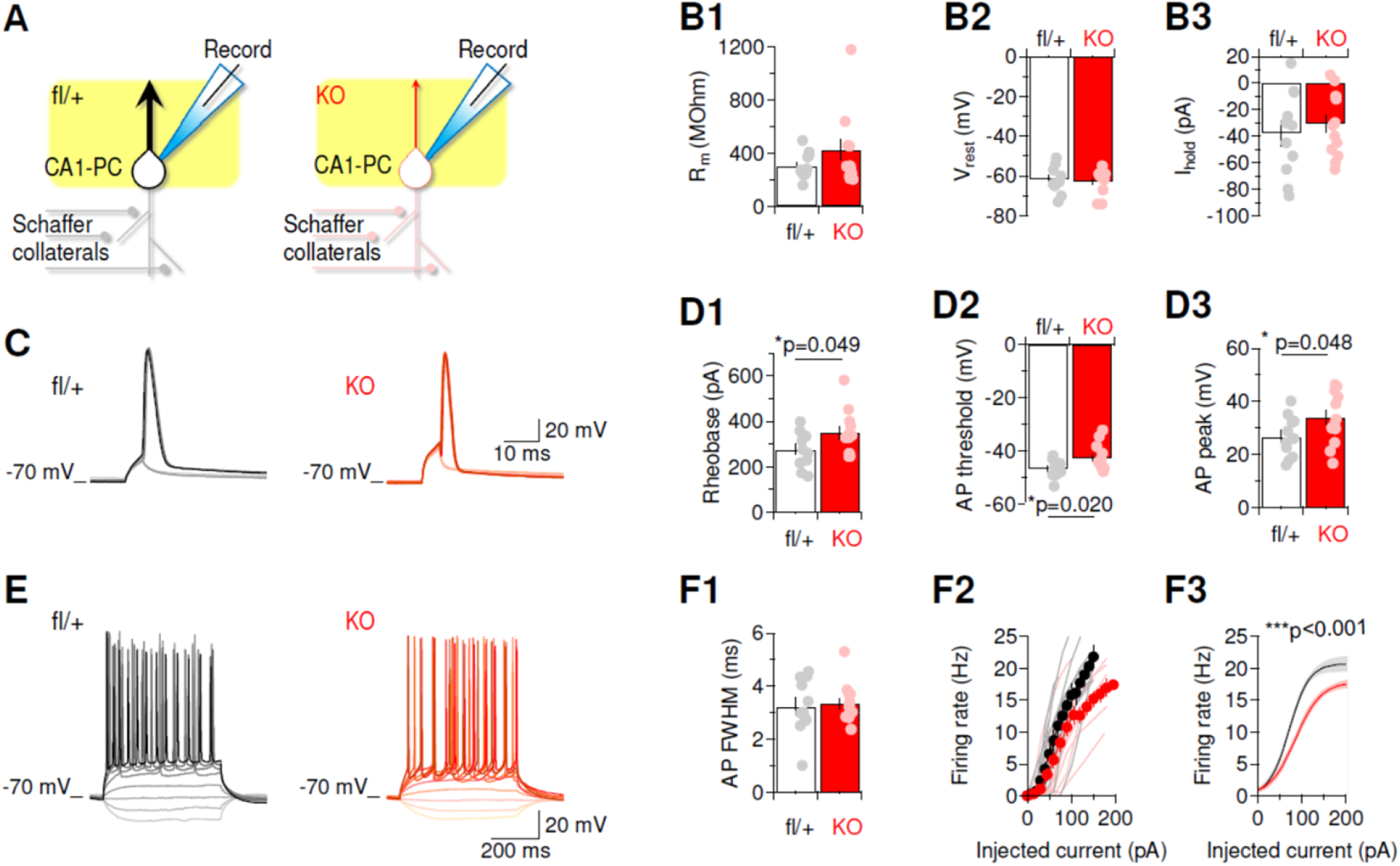
Passive and active membrane properties of hippocampal CA1 pyramidal cells in control and bLRRC8A KO mice. (A) Schematic representation of the experimental design and results. Note the reduced excitatory output in bLRRC8A KO hippocampus (red arrow) compared to control cells (black arrow). (B) Average membrane resistance (B1), resting membrane potential (B2), and holding currents needed to maintain CA1-PCs at −70 mV (B3) in CA1-PCs from control (fl/+ and fl/fl, n=9) and KO mice (n=13). (C) Representative sub- and supra-threshold responses to 5 ms somatic current injections of increasing amplitude. (D) Average rheobase (D1), action potential threshold (D2), and action potential peak amplitude (D3) in CA1-PCs from control (n=11) and KO mice (n=13). (E) Representative voltage recordings in response to 500-ms positive and negative current step injections. (F1) Average of the action potential full width at half maximum (FWHM) in CA1-PCs from control (n=11) and KO mice (n=13). (F2) Firing rate vs. current (f/I) plots showing the firing rates evoked by somatic current step injections of different amplitudes. Raw data for control slices are shown in grey. Raw data for KO are shown in pink. The symbols represent the average of all data collected from control (black, n=12) and KO mice (red, n=13). (F3) Sigmoidal fits of the data in F2. Data represent mean ±SEM, analyzed using unpaired t-test.

To determine how these changes in cell excitability affect sustained firing of CA1-PCs, we applied longer (500 ms) supra-threshold current steps (Fig. 5E). The firing rate-current plots showed that the firing rate of CA1-PCs in bLRRC8A KO mice was smaller than in control (fl/+ and fl/fl) mice over a range of positive current injections, leading to a rightward shift in the f/I plots (Fig. 5F2-F3). Together, these results suggest that loss of LRRC8A protein and VRAC activity impairs the excitability of CA1-PCs, with potential consequences on information flow in and out of the hippocampus.

### 3.6 LRRC8A bKO modifies GABAergic inhibition onto CA1 pyramidal cells

The propensity of CA1-PCs to generate action potentials in response to incoming stimuli depends on their ability to integrate in space and time both excitatory and inhibitory synaptic inputs. Therefore, we performed experiments to identify changes in excitatory and inhibitory synaptic transmission in the CA1 region of LRRC8A bKO mice. We first performed extracellular field recordings from the stratum radiatum. Here, a recording electrode and a bipolar stimulating electrode were both placed in the stratum radiatum, 100-150 μm away from each other (see Methods; Fig. 6A). Schaffer collateral stimulation evoked a complex voltage waveform, composed of a fast-rising and fast-decaying fiber volley (FV), followed by a slow-decaying field post-synaptic potential (fPSP; Fig. 6B). The FV amplitude provides a proxy readout of the number of activated afferent fibers. Therefore, by measuring the ratio between the fPSP slope and the FV amplitude, we measured the magnitude of the post-synaptic response for a given number of activated afferent fibers. This ratio was similar in control and KO groups (Fig. 6B). Blocking GABA_A_ receptors with picrotoxin (PTX, 100 μM) caused a significant increase in the fEPSP/FV ratio in control (fl/+ and fl/fl) and KO mice. The effect of picrotoxin was similar across these genotypes (Fig. 6C). Together, these results suggest that LRRC8A deletion produces no overt change in the input/output ratio at Schaffer collateral synapses and in neuronal sensitivity to GABAergic inhibition (but see the detailed analysis below).

**FIGURE 6.**
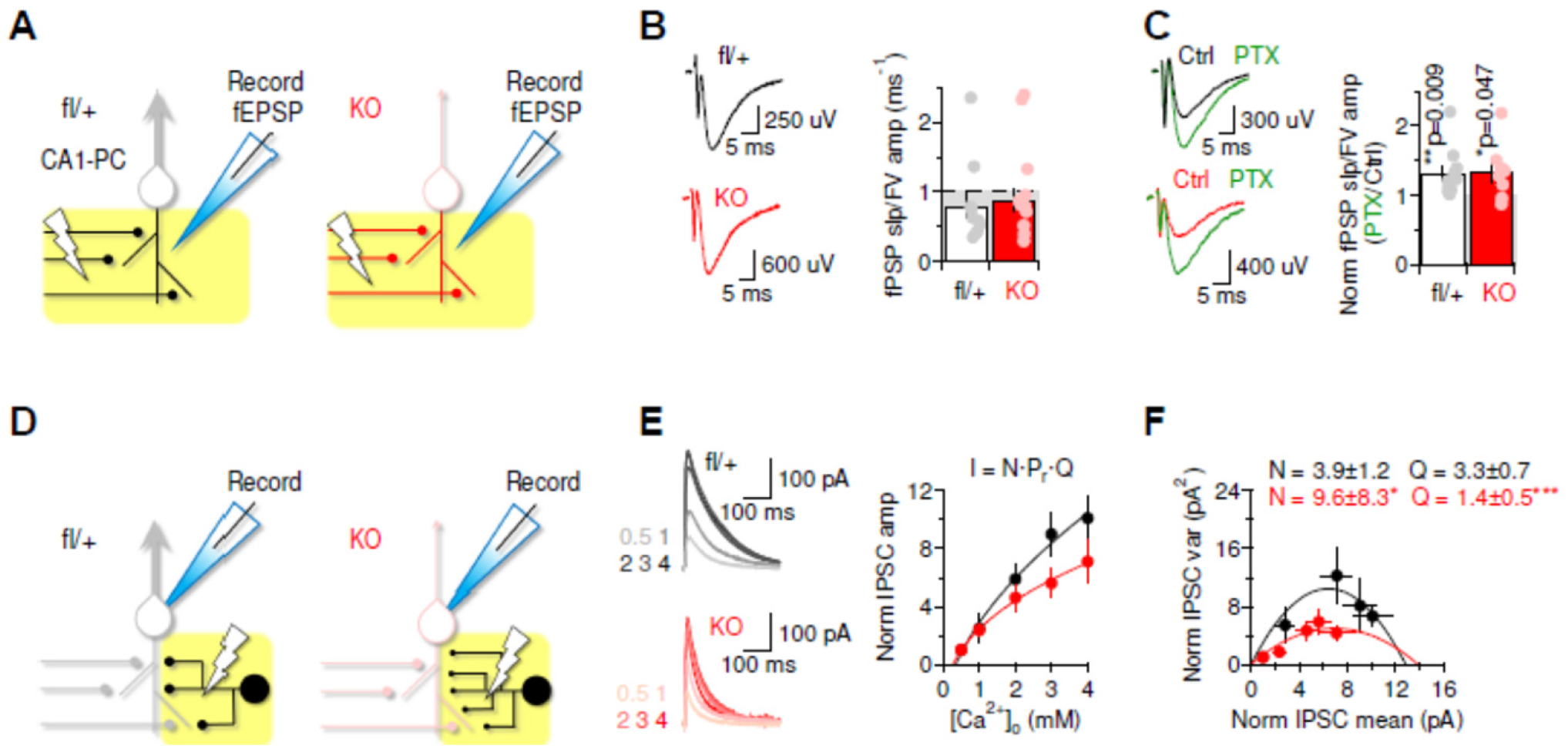
Effect of LRRC8A deletion on CA1 field recordings and quantal properties of GABAergic inhibition onto CA1 pyramidal cells. (A) Schematic representation of experimental design and results of extracellular field excitatory post-synaptic potential (fEPSPs) recordings in response to electrical stimulation of Schaffer collaterals, highlighted in yellow. (B) Representative field recordings from control (fl/+) and KO slices (left panels). Each trace is the average of 20 consecutive recordings. The summary bar graph shows the ratio between the fEPSP slope and the fiber volley (FV) amplitude in control (n=11) and KO mice (n=16; right panel). (C) The GABA_A_ receptor competitive antagonist picrotoxin (PTX, 100 μM) has a similar effect on the extracellular field recordings in control (n=11) and KO mice (n=8). The p values refer to in-slice comparisons of the input/output ratio before and after adding PTX. (D) Schematic representation of experimental design and results of whole-cell recordings of CA1 pyramidal cells in response to IPSCs from inhibitory interneurons, highlighted in yellow. The increased branching and smaller size of inhibitory connections refer to the larger N and smaller Q values measured in KO mice. (E) Representative IPSCs recorded at different extracellular calcium concentrations in control and KO slices (left panels). Relationship between [Ca^2+^]_o_ and the IPSC amplitude, normalized to its value at [Ca^2+^]_o_= 0.5 mM in control and KO slices (right panel). (F) The results of variance-mean analysis. The data are shown as normalized by the values obtained at [Ca^2+^]_o_= 0.5 mM. The solid curves are the parabolic fit of the data (see Methods). The estimates of N and Q are reported at the top. Data are shown as mean ±SEM, analyzed by unpaired t-test (B, C).

The latter results do not rule out the possibility that more subtle changes in GABAergic inhibition may occur at inhibitory synapses onto CA1-PCs. For this reason, we performed whole-cell recordings in voltage-clamp mode to record inhibitory postsynaptic currents (IPSCs) from CA1-PCs held at 0 mV (the estimated reversal potential for glutamatergic currents). At this potential, GABAergic IPSCs can be isolated pharmacologically by adding the AMPA receptor antagonist NBQX (10 μM) and the NMDA receptor antagonist APV (50 μM) to the extracellular solution, as outward currents (Fig. 6D). We recorded IPSCs at a range of extracellular calcium concentrations (0.5-4 mM), to vary release probability. The IPSC amplitude increased exponentially when increasing the extracellular calcium concentration, consistent with previous work.^54^ This relationship, however, differed between control (fl/+ and fl/fl) and KO slices (Fig. 6E). This effect is unlikely to be accounted for by changes in release probability, because it was not associated with significant changes in the IPSC paired-pulse ratio ([Ca^2+^]_o_=2 mM PPR_control_: 0.88±0.07 (n=7), PPR_KO_: 1.05±0.08 (n=14), p=0.12). We can gain insights about the quantal parameters *N* (the number of release sites), and *Q* (the quantal size) by analyzing the trial-to-trial fluctuations in the IPSC amplitude, using multiple-probability fluctuation analysis (MPFA). ^42–44^ One of the main strengths of the MPFA analysis is that it does not rely on assumptions of uniform release probability distribution across release sites, which might occur at hippocampal inhibitory synapses. By using the MPFA, we identified two major changes in KO mice, due to a significant increase in *N* and a decrease in *Q* (Fig. 6F). This means that there are more functional GABAergic synapses onto CA1 pyramidal cells in KO mice, but each connection is weaker because it evokes a smaller current in response to the release of a synaptic vesicle. These effects can compensate each other and can potentially account for the lack of overt changes detected using field recordings. Together, these results suggest that there are notable functional changes in the properties of GABA release from inhibitory neurons onto CA1-PCs in KO animals, which cannot be detected using extracellular field recordings (for functional implications see Discussion).

### 3.7 LRRC8A bKO results in decreased GAT-1 levels but no significant changes in other GABAergic cell markers

To determine whether there was any dysfunction in GABA synthesis, vesicular loading of GABA, or neurotransmitter reuptake, changes in protein expression of glutamic acid decarboxylase isoforms (GAD65/67), vesicular GABA transporter (VGAT), and the major plasmalemmal GABA transporter isoform (GAT-1) were examined. The semi-quantitative western blotting showed no differences in the expression of GAD65/67 or VGAT proteins in whole-brain protein lysates (Fig. 7B, C, full blots in Suppl. Fig. 7.1 and 7.2). Protein levels of GAT-1 in LRRC8A bKO brains were reduced by 38%, (p<0.01, Fig. 7D, full blots in Suppl. Fig. 7.3). The GAT-1 transporter is predominantly located presynaptically in interneurons but is also expressed in astrocytes.^55;56^ To explore potential cell loss in individual interneuron populations, we immunolabeled and counted parvalbumin-positive cells in the CA1 pyramidal cell layer (Fig. 7F1) and in the cerebral cortex above the hippocampus in the same slices (Fig. 7E1). Quantitative analysis revealed no significant changes in PV-positive cell populations in either of the analyzed brain regions (Fig. 7E2,F2).

**FIGURE 7.**
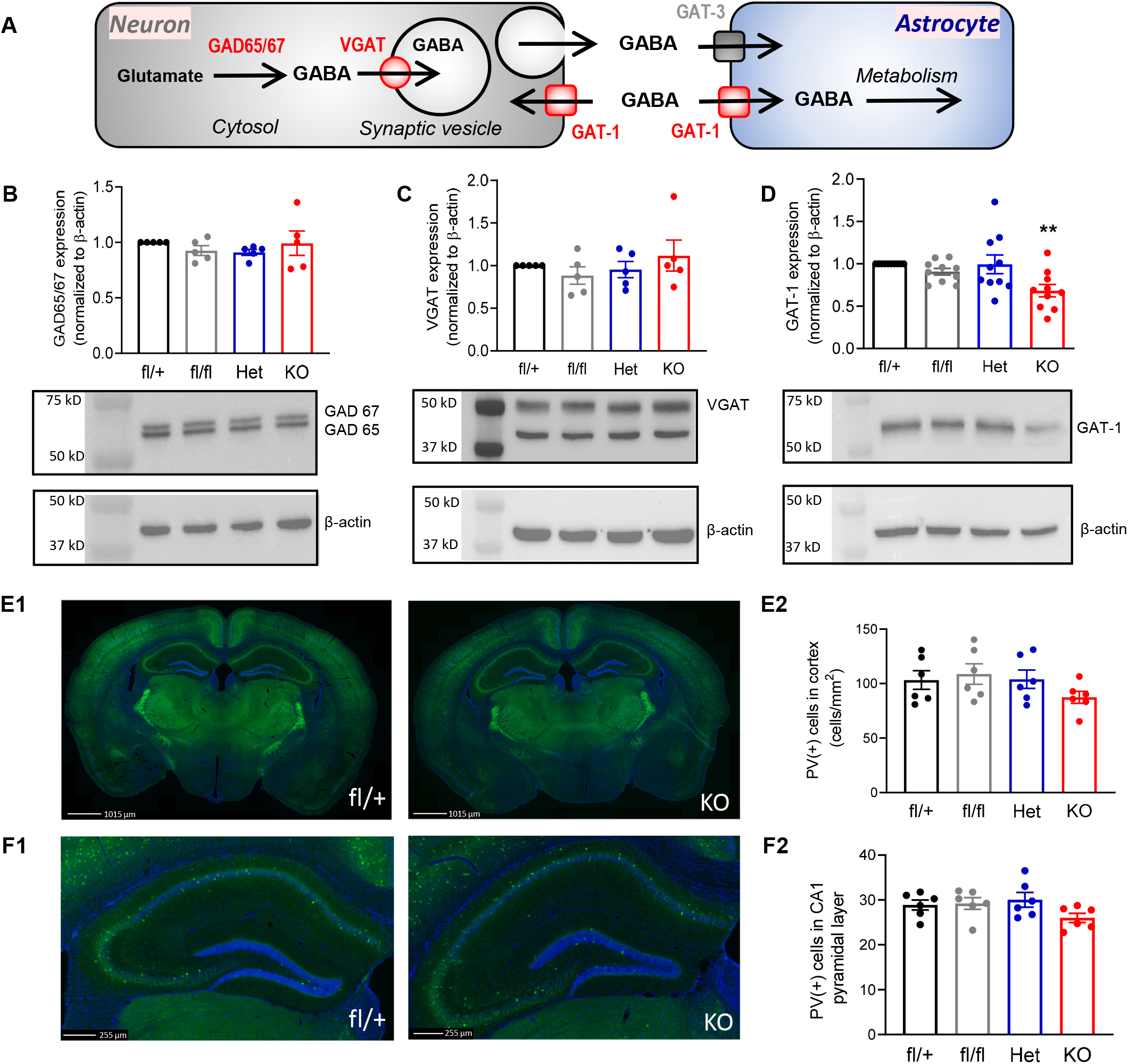
Expression of proteins involved in GABA synthesis, vesicular packaging, or re-uptake in whole-brain lysates and numbers of parvalbumin-positive interneurons in bLRRC8A KO mice. (A) Schematic of the role of glutamate decarboxylase (GAD65/67), vesicular glutamate transporter (VGAT), and membrane GABA transporters (GAT-1 and GAT-3) in GABAergic neurons and astrocytes. (B-D) Western blot analysis of (B) GAD65/67, (C) VGAT, or (D) GAT-1 protein levels in whole-brain lysates. Insets show representative western blot images for each protein and matching β-actin loading controls on the same membrane. Data are the mean values ±SEM of immunoreactivity in 5 brain or 10 (in D) lysates per group normalized to β-actin loading controls and then to fl/+ control in each set. **p<0.01, fl/+ vs. KO in D, Student’s t-test with Bonferroni correction. (E1) Representative immunohistochemistry images of parvalbumin-positive (green) and DAPI (blue) stained brain slices in fl/+ controls (left) and KO (right) brains. (E2) Quantification of parvalbumin-positive cells in medial cortex, in the region identified by the dotted white line. Data are the mean values ±SEM of 6 animals per genotype. (F1) Magnified images and quantification of the parvalbumin immunohistochemistry staining in the hippocampus on animals from E1. (F2) Quantification of parvalbumin-positive cells in CA1 pyramidal layer. Data are the mean values ±SEM of 6 animals per genotype. Analyzed by one-way ANOVA with Bonferroni correction.

### 3.8 Downregulation of glutamate transporter GLT-1 and modifications in hippocampal glutamine synthetase and neurotransmitter levels in LRRC8 KO brains

Because we found changes in astrocytic GFAP immunoreactivity, we next quantified the expression levels of proteins involved in astrocytic transport and synthesis of the GABA precursor molecules, glutamate and glutamine. As a critical step in astrocyte-neuron metabolic coupling, extracellular glutamate is taken up into astrocytes through glutamate transporters, including GLT-1 (Fig. 8A). Western blot analysis revealed that whole-brain expression of GLT-1 in bLRRC8A KO mice was reduced, on average by 40% but even more dramatically in some brains (p<0.05; Fig. 8B, full blots in Suppl. Fig. 8.1). Astrocytes convert glutamate to glutamine via the activity of glutamine synthetase and further export glutamine to neurons for production of glutamate and GABA (see Fig. 8A and review^57^). Whole-brain lysates showed a 14% reduction in the level of the glutamine-producing glutamine synthetase in bLRRC8A KO brains (p<0.05, Fig 8C, full blots in Suppl. Fig. 8.2). To investigate whether there were more local changes in the glutamine synthetase expression, we performed immunohistochemical analysis of this enzyme in the hippocampal CA1 stratum radiatum, the region probed in our electrophysiology recordings, and additionally stratum lacunosum moleculare, and stratum moleculare (Fig. 9). As compared to fl/+ and fl/fl controls, in these hippocampal regions, the integral glutamine synthetase fluorescence intensity was consistently reduced by ∼ 17%, 24%, and 19%, respectively, but these changes were not statistically significant after multiple comparisons (Fig. 9B1-B3). The apparent decrease in glutamine synthetase expression can be clearly seen in representative high-magnification confocal images of the stratum radiatum (compare Fig. 9A3 and A4). Slices were co-stained with GFAP to confirm astrocytic localization, and reactive gliosis in LRRC8A bKO animals (Fig. 9A3 and A4). We recapitulated strong and statistically significant increases in integral GFAP immunoreactivity in all three analyzed regions of the hippocampus (Fig. 9C1-C3).

**FIGURE 8.**
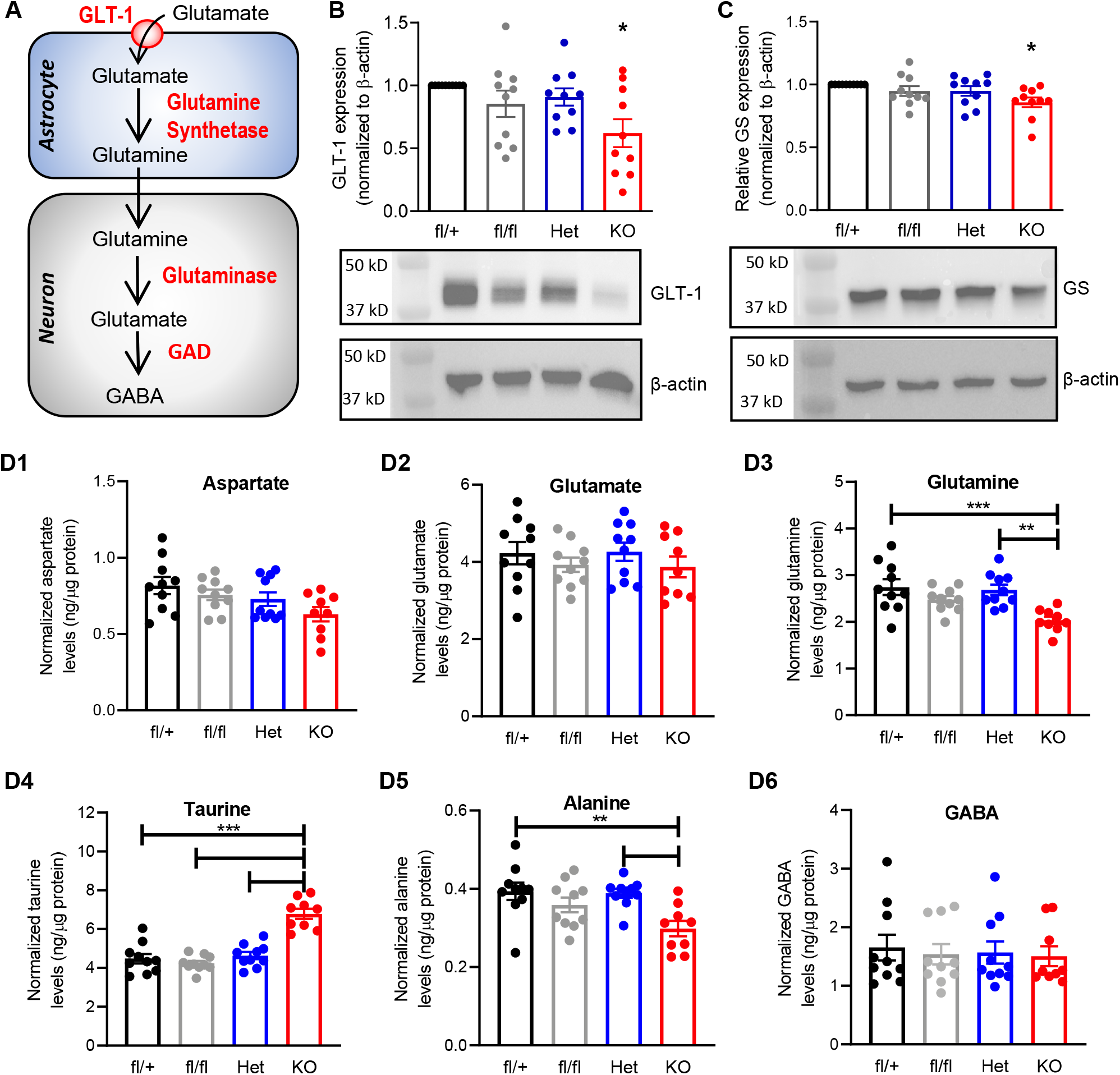
Effect of LRRC8A deletion on glutamate transporter and glutamine synthetase levels, and amino acid content in the brain tissue. (A) Schematic of glutamate-glutamine-GABA cycle in astrocytes and neurons. (B-C) Western blots analysis of (B) the glutamate transporter GLT-1 and (C) glutamine synthetase levels in whole-brain lysates, normalized to β-actin loading controls and then to the fl/+ control in each set. Data are the mean values ±SEM of 10 protein lysates for each genotype. Insets show representative western blot images for GLT-1 or glutamine synthetase and β-actin re-probed on the same membrane. *p < 0.05, KO vs. fl/+, one-sample t-test with Bonferroni correction. (D) HPLC analysis of the tissue levels of aspartate (D1), glutamate (D2), glutamine (D3), taurine (D4), alanine (D5), or GABA (D6) in the whole-brain lysates from four genotypes, normalized to protein levels. Data are the mean values ±SEM of 9-10 independent animals per genotype, ** p <0.01, ***p<0.001, Kruskal-Wallis with Dunn’s test for multiple comparisons (D1), or one-way ANOVA with Bonferroni correction (D2-D6).

**FIGURE 9.**
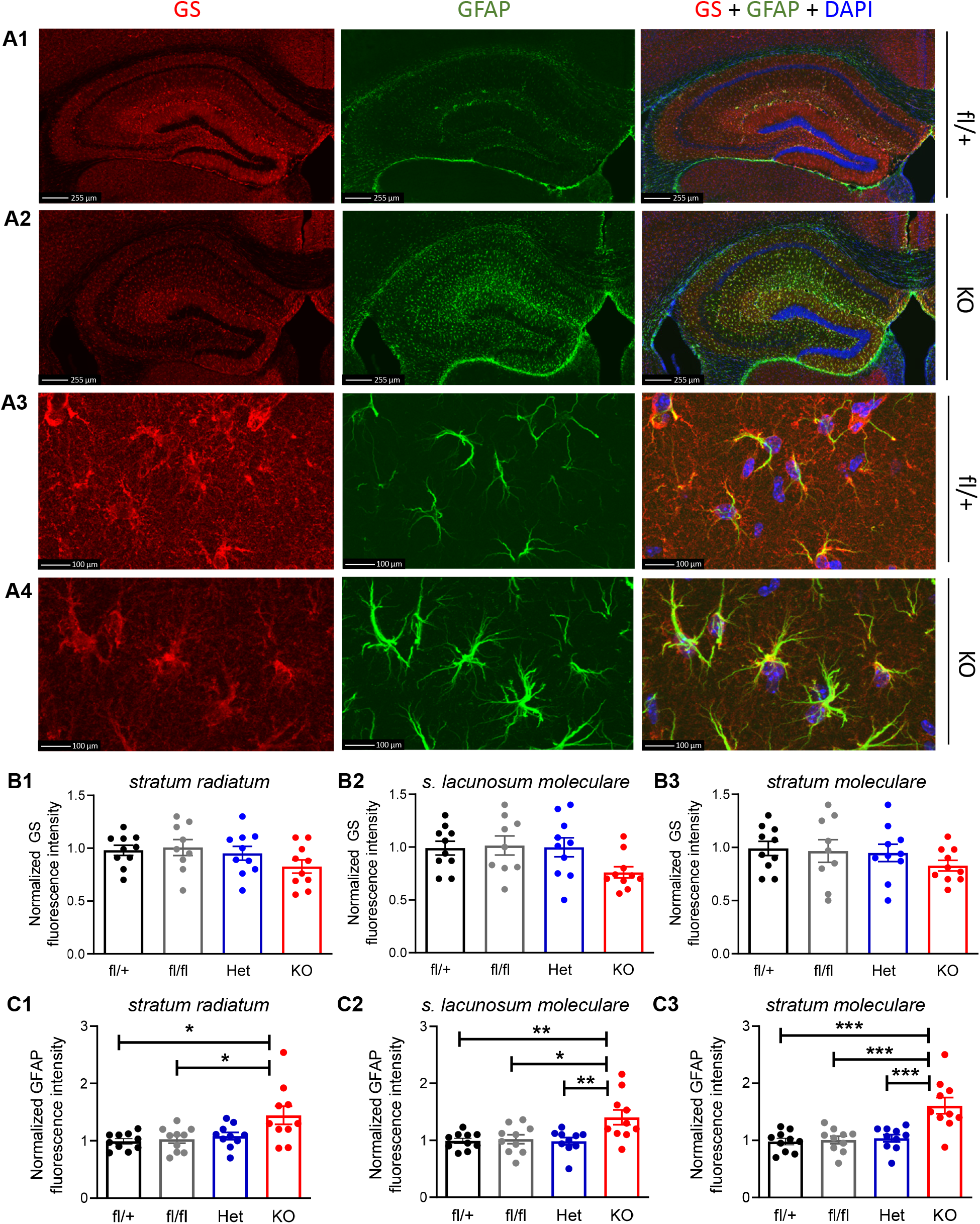
Immunohistochemical analysis of glutamine synthetase expression in the hippocampus following LRRC8A deletion. (A) Representative images of the immunoreactivity for glutamine synthetase in the entire hippocampus (A1, A2) and high-magnification images of the CA1 *stratum radiatum* (A3, A4) in fl/+ controls and KO brains. Left panel, glutamine synthetase (GS) immunoreactivity (red). Middle panel, astrocytic GFAP expression (green). Right panel, overlay of glutamine synthetase and GFAP immunoreactivity in the same brain slices, counterstained with DAPI (blue). White box indicates the region of high-magnification images (A3, A4). (B) Relative glutamine synthetase immunofluorescence in the CA1 *stratum radiatum* (B1), *stratum lacunosum moleculare* (B2), and *stratum moleculare* (B3) in four genotypes. (C) Relative GFAP signal in the CA1 *stratum radiatum* (C1), *stratum lacunosum moleculare* (C2), and *stratum moleculare* (C3) in four genotypes. Data are the mean values ±SEM of 10 animals per group, normalized to the average fluorescence intensity of each set. Analyzed by two-way ANOVA with a fixed effect of genotype and a random effect of staining set number to remove between-set variability. *p<0.05, **p<0.01, ***p<0.001, as indicated.

Consistent with the idea of disruption in the glutamine synthesis pathway, we found statistically significant reductions in the whole-brain levels of glutamine in KO mice, as determined by HPLC analysis and compared to fl/+ controls and Het animals (Fig. 8D3). There was also a significant reduction in the whole-brain levels of alanine (Fig. 8D5). In the brain, alanine synthesis and transport are known to be connected to processes involving both GABAergic and glutamatergic signaling via transamination and the lactate-alanine shuttle.^58;59^ There were no statistically significant changes in the levels of aspartate (Fig. 8D1), glutamate (Fig. 8D2), or GABA (Fig. 8D6), but this may be due to homeostatic regulation or experimental variability. In striking contrast to changes in other amino acid levels, LRRC8A deletion caused a significant increase (∼50%) in the brain levels of taurine (Fig. 8D4, and Suppl. Fig. 8.3). To the best of our knowledge, there is no metabolic connection between production of taurine and GABA, glutamate, and glutamine. However, because taurine acts as an endogenous agonist at GABA_A_ and glycine receptors^60;61^, its increase may be compensatory to limit hyperexcitability in LRRC8A bKO brains. Overall, these data suggest reduced astrocytic uptake of glutamate and diminished export of glutamine as a potential contributor to the decreased quantal size of GABAergic inputs and perhaps seizure development observed in LRRC8Ab KO mice (see Discussion).

## 4 DISCUSSION

The most striking observation of our work is the 100% lethality and seizure phenotype in early to late adolescence in mice with the brain-specific deletion of the LRRC8A protein. None of the 32 group-housed LRRC8A bKO animals, which were not utilized for histological and biochemical analyses, survived past the end of postnatal week nine. The same pattern of mortality of the *Nestin*^Cre^-driven LRRC8A deletion was recently reported by another group, who utilized this mouse in stroke studies but did not explore the cause of death.^62^ The severity of the brain-wide LRRC8A KO phenotype loosely resembles the dramatic embryonic and early postnatal mortality of mice with global deletion of LRRC8A, who die due to severe defects in multiple organs.^22^ However, our brain-specific LRRC8A KO mice developed without apparent organ abnormalities, in the brain or elsewhere.

We did not expect brain LRRC8A KO to be lethal for the following reasons. The recent work pursuing deletion of astrocytic LRRC8A with the relatively selective *mGFAP*^Cre^ promoter produced completely viable animals with marginally reduced neuronal excitability and high resilience against ischemic brain damage.^35^ These reported findings of a “less excitable” brain are consistent with the long-standing idea that astroglial VRAC promotes neuronal excitability and brain damage via physiological and pathological release of the excitatory amino acid transmitters, aspartate and glutamate.^9;30–32^ Since the *Nestin*^Cre^-driven LRRC8A deletion targets three major types of brain cells, we anticipated a more pronounced neurological phenotype and perhaps developmental defects. Yet, LRRC8A bKO mice developed without gross abnormalities, including no changes in numbers of neurons and astrocytes, or myelination by oligodendroglia. Counterintuitive to the idea of “less excitable” brain, all LRRC8A bKO mice (among those who were systematically investigated) developed recurrent spontaneous seizures. These seizures represent the most likely cause of mortality, because four out of six continuously monitored bLRRC8A KO mice died with a generalized seizure, which preceded the moment of death. Among the two remaining bLRRC8A KO animals with spontaneous seizures, one died without coinciding generalized seizure and another unexpectedly lived for more than six months and was euthanized. A subset of the observed seizures, including all of those that preceded death, were of a brainstem-tonic manifestation, suggesting that ictal discharges can and do propagate to the brainstem regions important for life, such as respiratory and cardiac centers. Anomalous activity in the brainstem has been implicated in sudden unexpected death in human epilepsy and mouse models (e.g.,^63^), including in our recent model work on the spontaneous seizures induced by short-term flurothyl treatment.^38^

Pinpointing the cellular and molecular mechanism(s) for changes in neuronal excitation proved to be difficult. We started by measuring neuronal excitability in the CA1 region of the hippocampus. The whole-cell recordings in CA1 pyramidal neurons revealed modest reductions in their excitability, resembling the recent findings in astrocytic LRRC8A KO mice.^35^ This effect on its own is unlikely to account for the onset of seizures. Therefore, in the search for pathological mechanism, we further probed for changes in synaptic transmission. The recordings of field potentials evoked in stratum radiatum by the stimulation of Schaffer collaterals did not identify major differences in the excitatory responses, or in the sensitivity of field potentials to the blockade of GABA_A_ receptors. However, this general lack of effect of GABA blockers, does not completely rule out the possibility of changes in inhibitory transmission. In fact, MPFA analysis in our whole-cell recordings in the CA1 pyramidal cells indicated that there was a significant increase in the number of functional inhibitory connections in bLRRC8A KO mice coupled with a marked reduction in the quantal size of GABAergic IPSCs. Existing literature strongly suggests that changes in GABAergic circuitry can increase predisposition to seizures and promote epileptogenesis (e.g.,^64;65^ and reviews^66;67^). Our molecular examination of GABAergic machinery found no alterations in the whole-brain levels of the GABA-producing enzymes, GAD65 and GAD67, or the vesicular GABA transporter VGAT in bLRRC8A KO brains. However, we did find a 38% reduction in the levels of the major plasmalemmal GABA transporter GAT-1. This latter discovery may be instructive on two levels. (i) Human and animal studies link partial or complete loss of GAT-1 (*SLC6A1*) activity to epileptogenesis.^68–70^ (ii) GAD-1 KO mice demonstrate dramatically reduced quantal size of GABA release.^71^ Yet, the similarity between bLRRC8A KO and loss-of-function GAT-1 phenotypes is only partial. GAT-1 deletion and loss-of-function GAT-1 mutations do not produce distinct mortality phenotypes and are largely associated with epilepsies of the myoclonic atonic type and reduced mobility.^69;70^ Therefore, bLRRC8A KO deficits likely involve additional mechanism(s).

In further search of the molecular changes responsible for modified GABAergic signaling, we turned to the brain glutamine-glutamate/GABA cycle. In the CNS, both glutamatergic and GABAergic neurons use astrocyte-derived glutamine for the synthesis of glutamate and GABA, respectively.^59;72^ Production of GABA consumes between 10-25% of glutamine flow^73;74^, and is highly sensitive to inhibition of any step of the glutamine-glutamate cycle.^75;76^ The bulk of astrocytic glutamine is, in turn, produced via recycling the neuronal glutamate taken up via the high affinity astrocytic glutamate transporter GLT-1.^77^ In this context, we found very strong (albeit somewhat variable) decreases in the levels of GLT-1 in LRRC8A bKO. To compare, global deletion of GLT-1in mice produces spontaneous lethal seizures with the onset of mortality at postnatal week 3 and 50% survival by week 6.^78^. Much like in our study, GLT-1 KO deaths are not preceded by noticeable health problems but rather coincide with the onset of spontaneous seizures.^78^ Along the same line, postnatal astrocytic deletion of GLT-1 using *Gfap*^CreERT2^ produces spontaneous seizures and mortality with the onset at 4-6 week and the median survival at 23 weeks of age.^79^ Other relevant changes we identified in LRRC8A bKO CNS were reductions in total and hippocampal glutamine synthetase immunoreactivity and a matching drop in the total brain glutamine content. Numerous reports link the loss or inhibition of astrocytic glutamine synthetase to hyperexcitation and the development of low-grade status epilepticus^80–82^, and propose the loss of the glutamine synthetase protein as a potential cause of human temporal lobe epilepsy.^83^ One mouse study linked the induction of reactive gliosis, to downregulation of glutamine synthase and the deficit in inhibitory, but not excitatory synaptic function.^84^ A model of kainate-induced chronic epileptogenesis produces very similar to our findings moderate decreases in glutamine synthesis, with no matching decreases in whole-brain GABA levels.^85^ Another electrophysiology study has reported that pharmacological inhibition of glutamine synthetase decreases glutamine import, triggers progressive loss of GABAergic drive and reduces quantal GABA release at individual GABAergic synapses in hippocampal slices.^75^ It is tempting to draw a parallel between these published results and our own MPFA data and conclude that inhibition of glutamine export, due to loss of glutamine synthetase and/or deficiency of GLT-1, is responsible for the reduced GABAergic tone and animal seizures/mortality. The maturation of the glutamine-glutamate cycle in mice happens postnatally and is completed by, or after, week four^74;86^, which precedes the onset of mortality in bLRRC8A KO animals.

A complementary theoretical mechanism that can contribute to hyperexcitability in LRRC8A bKO mice is the failure of dynamic control of cellular volume and extracellular space. During development, the volume of extracellular space in the brain is reduced progressively and significantly to an average value of ∼20% of the total brain volume in adulthood (reviewed in^87^). However, the volume of brain interstitial space does not stay constant and is dynamically reduced during neuronal excitation, mostly due to the swelling of astrocytes which accumulate ions and neurotransmitters (e.g.,^88–91^ and reviews^33;87^). Deletion of LRRC8A-containing VRAC results in the loss of cell volume regulation, including in brain cells^11;12;92^, and may blunt homeostatic control of the extracellular space. The persistent cellular swelling during periods of enhanced neuronal activity is sufficient to increase the extracellular K^+^ and excitatory neurotransmitter levels, and cause hyperexcitation.^33;93^ In fact, such swelling has been proposed as an important driving cause in epileptogenesis.^33;93–96^ The human disease equivalent of reduced VRAC activity in the brain is megalencephalic leukoencephalopathy with subcortical cysts (MLC). MLC is a childhood-onset disorder caused by mutations in the astrocyte-specific protein MLC1, and the associated GlialCAM, and is characterized by the loss of white mater and vacuolation in myelin and astrocytes.^97–99^ MLC causes disruptions in ion and water homeostasis which are thought to develop due to reduced activity of the LRRC8A-containing VRAC ^99–101^. The majority of MLC patients have spontaneous seizures.^102;103^ A study in MLC1-null and GlialCAM-null animals found spontaneous interictal activity, low seizure threshold and increased seizure severity in kainate-treated animals, along with impaired extracellular K^+^ clearance. ^103^ The authors of the latter work explained animal hyperexcitability by ineffective K^+^ clarence in astrocytic networks with reduced VRAC activity. Although the latter mechanism is difficult to probe in brain slice preparations, it may also contribute to spontaneous seizures in bLRRC8A KO mice.

In conclusion, we established that the LRRC8A-containing volume-regulated anion channels are dispensable for normal brain development but contribute critically to the control of brain tissue excitability in adolescence. Mice with the brain-wide deletion of the essential VRAC subunit, LRRC8A, in neurons, astrocytes and oligodendrocytes, develop a seizure phenotype and die in early-to-late adolescence. We identified a number of changes that are schematically summarized in Fig. 10 and may contribute to this lethal phenotype. These include modified GABAergic transmission, reactive astrogliosis, downregulation of the glutamate transporter GLT-1 and the GABA transporter GAT-1, and partial loss of astrocytic glutamine synthetase in the hippocampus and the whole brain. It is likely that the alterations in neurotransmitter transport and metabolism compound to produce increased brain excitability, spontaneous seizures, and animal death. Although the causal role for each individual enzyme and transporter in LRRC8A bKO outcomes has not been experimentally probed, it is noteworthy that every molecular change identified by us has been mechanistically linked to promoting epileptogenesis and animal mortality in other animal models. It remains to be established if reactive astrogliosis and disruptions in glutamate-glutamine cycle arise due to modifications in anion transport, autocrine and paracrine transmitter release, or intracellular signaling in VRAC-null cells.

**FIGURE 10.**
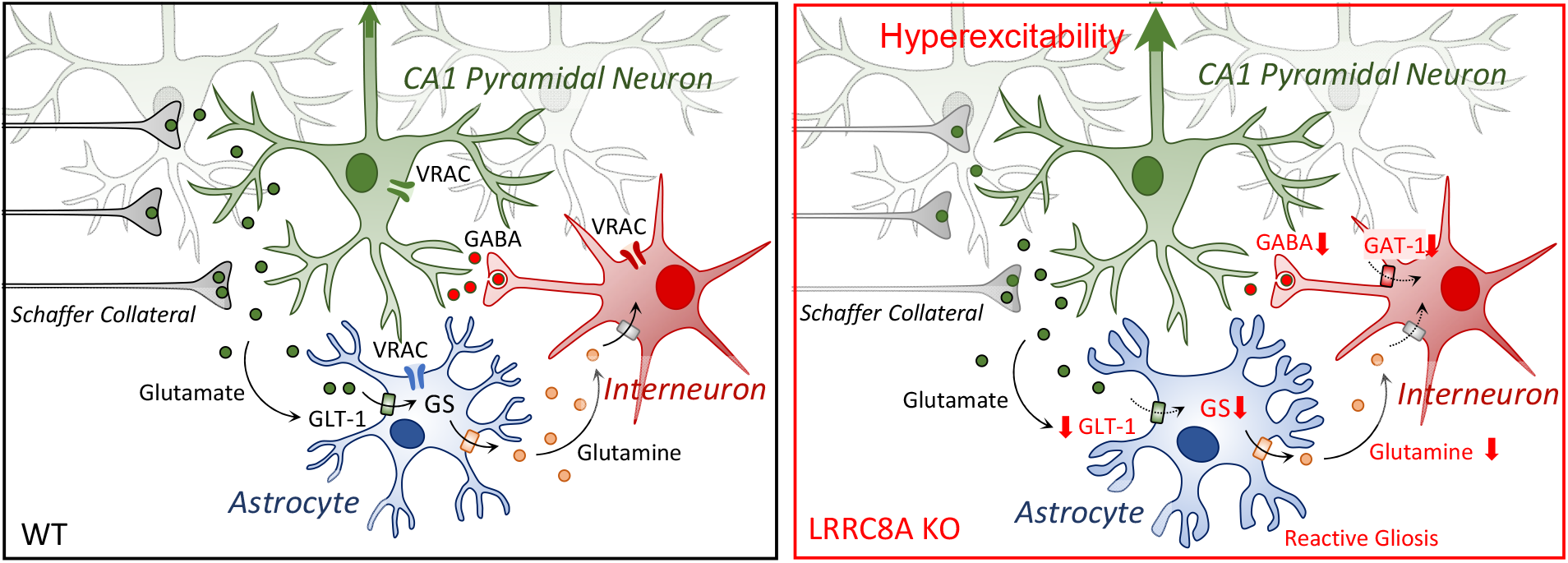
Graphical summary of the major findings in this work. Left: normal functional interactions between glutamatergic excitatory inputs (Schaffer collaterals, gray), glutamatergic pyramidal cells (green), GABAergic inhibitory interneurons (red), and astrocytes (blue) in a wild type (WT) hippocampus. Right: changes occurring upon removal of LRRC8A-containing VRAC, which likely lead to hyperexcitability. These include reductions in glutamate uptake (GLT-1), glutamine synthesis and supply (GS), glutamine levels, GABAergic inputs, and GABA reuptake (GAT-1).

## Acknowledgements

We thank Drs. Sophie Belin, Paul J. Feustel, Ashley Kopec, Joseph E. Mazurkiewicz, Yannick Poitelon, Damian S. Shin, Ling Wang, and Ms. Linda Barenboim for methodological advice on various aspects of this work.

## Author contributions

CSW and AAM conceived the study. CSW, PD, JWN, RJF, AS and AAM developed and finalized experimental design. CSW, PD, SO, JWN, RJF, and AS performed experiments and analyzed the data. RJF, YH, RS, AS, and AAM provided analytical tools and conceptual analysis. CSW, PD, JWN, RJF, AS, and AAM have written the manuscript. All authors contributed to the final manuscript edits.

## Preprint information

This manuscript has been deposited to bioRxiv in a preprint form under the Creative Commons license (CC-BY-NC-ND). doi: 10.1101/2020.05.22.109462

## Funding information

| Funding agency | Grant reference number | Author |
| --- | --- | --- |
| National Institute of Health | R01 NS111943 | Alexander A. Mongin |
| National Institute of Health | F31 NS111889 | Corinne S. Wilson |
| National Institute of Health | R01 NS092062 | Russell J. Ferland |
| National Institute of Health | R01 NS093045 | Yunfei Huang |
| National Science Foundation | IOS1655365, IOS2011998 | Annalisa Scimemi |

## Additional files

- Supplemental figures
- Media file of seizures in bLRRC8A KO mice

## Data availability

All primary data used this work and complete statistical analyses for each figure have been deposited to the Open Science Framework (accession number https://osf.io/3fwjc [registered on 2020/05/22] and the revision https://osf.io/puwz3 [registered on 2020/12/17] and https://osf.io/nauxg [registered on August 23, 2021]). Complete set of full-length western blot images, representative full-size immunohistochemistry images, and representative video recordings of mice with seizures are included in Supplemental Information and additionally available from the Open Science Framework.

