## Supplemental Figures for "Late adolescence mortality in mice with brain-specific deletion of the volume-regulated anion channel subunit LRRC8A"

C.S. Wilson *et al.* ● **SUPPLEMENTARY FIGURE 1.1**

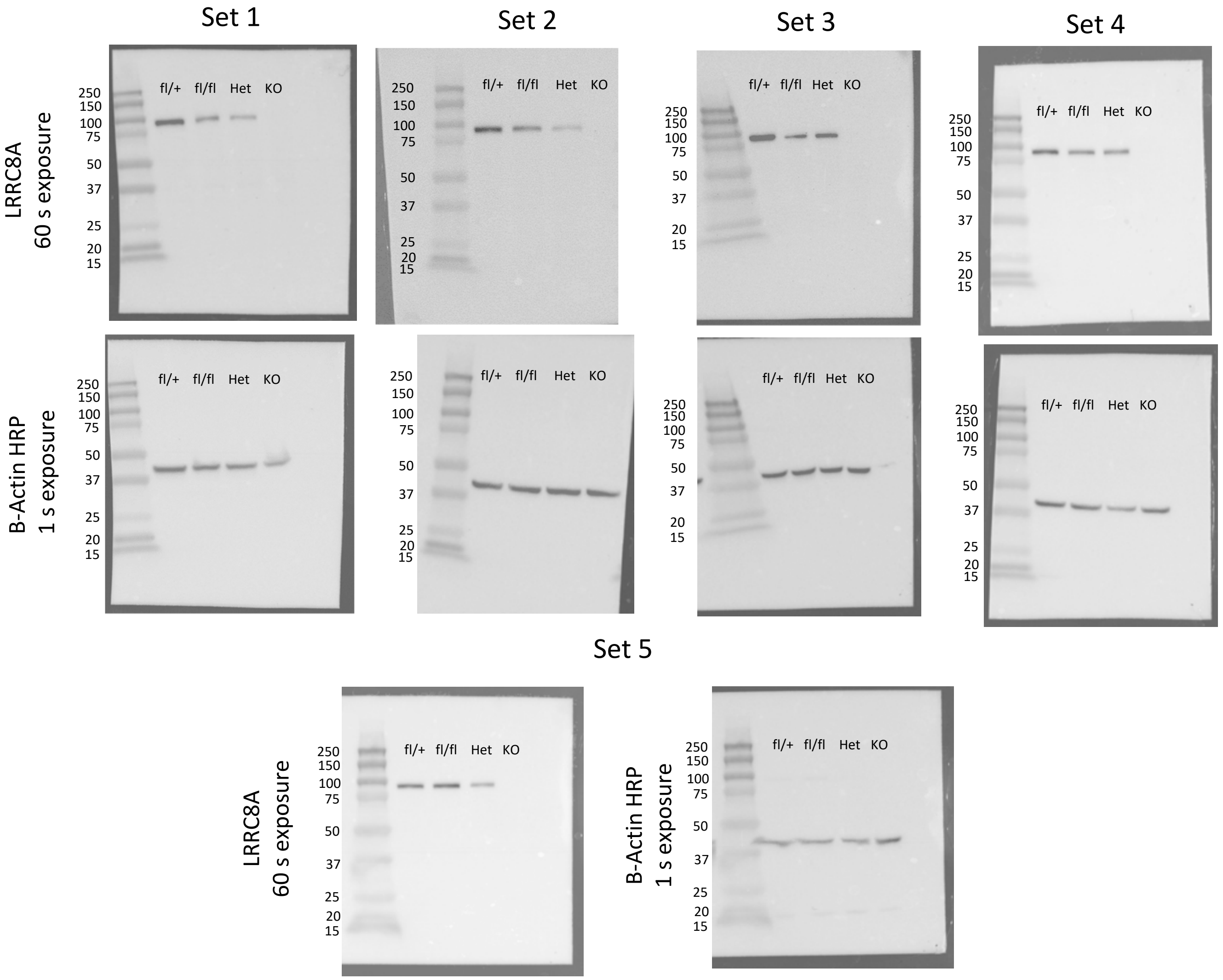


**Full-length western blots for LRRC8A protein and β-actin probed in five sets of whole-brain tissue lysates from fl/+, fl/fl, Het, or KO mice.** PAGE-separated proteins (20 µg/lane) were electrotransfered onto PDVF membrane, probed with monoclonal anti-LRRC8A antibody (Santa Cruz, cat #sc-517113, 1:500) and matching secondary antibody and visualized using an ECL kit and BioRad Gel Doc Imager. The membranes were then stripped, re-probed with the HRP-conjugated anti-β-actin antibody (Sigma Millipore, cat #A-3854 , 1:50,000), and processed in a similar manner.

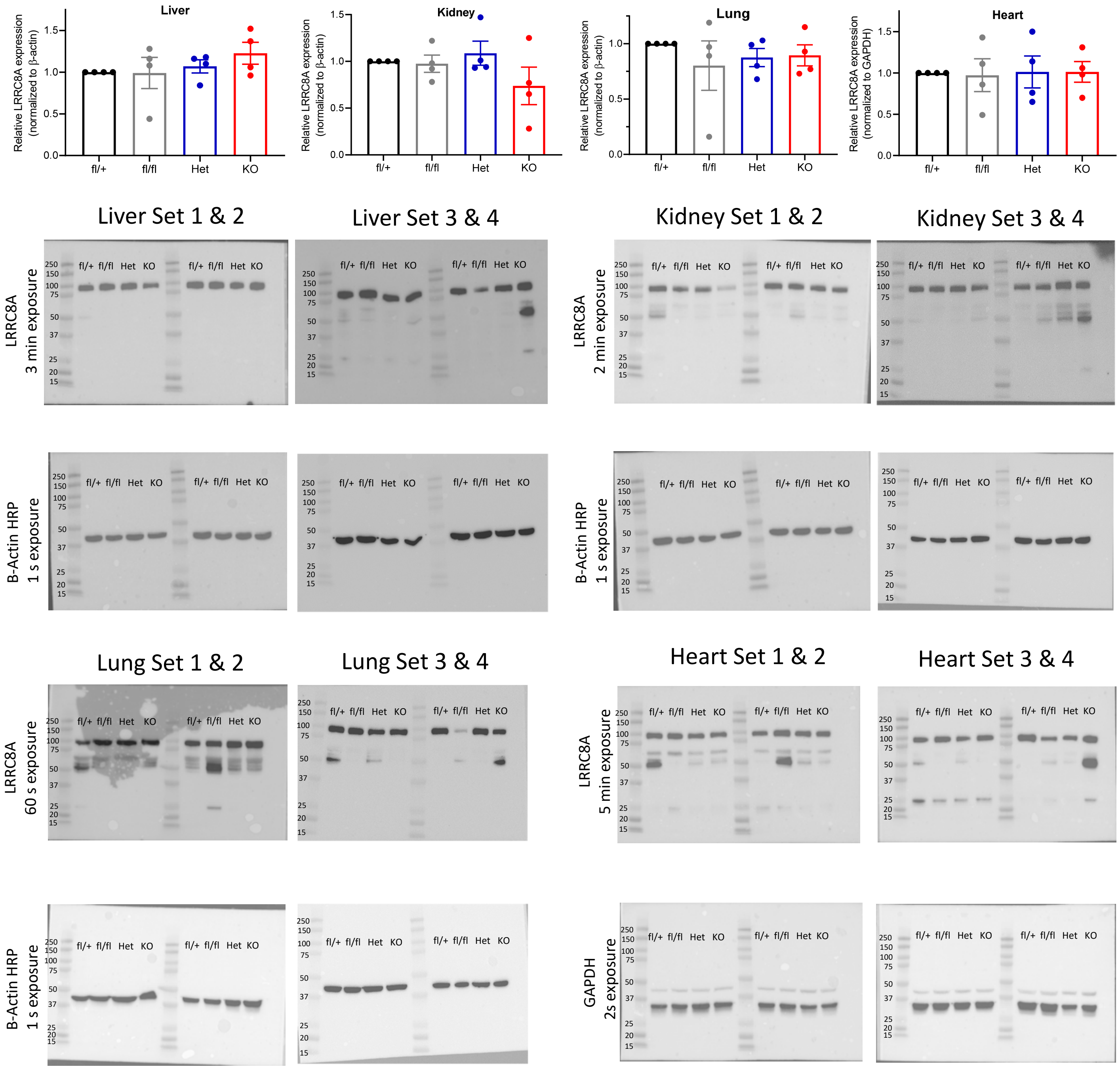


**Quantitative analysis and full-length western blots for LRRC8A protein and β-actin (or GAPDH) probed in the liver, kidney, lung and heart tissue lysates from four sets of fl/+, fl/fl, Het, or KO animals.** PAGE-separated proteins (20 µg/lane) were electrotransfered onto PDVF membrane, probed with monoclonal anti-LRRC8A antibody (Santa Cruz, cat #sc-517113, 1:500) and matching secondary antibody, and visualized using an ECL kit and BioRad Gel Doc Imager. The membranes were then stripped, re-probed with the HRP-conjugated anti-β-actin antibody (Sigma Millipore, cat #A-3854 , 1:50,000) and processed in a similar manner. In the heart tissue samples no expression of β-actin was detected; therefore, anti-GAPDH antibody (Millipore-Sigma, cat #G9545, 1:10,000) was used as a loading control.

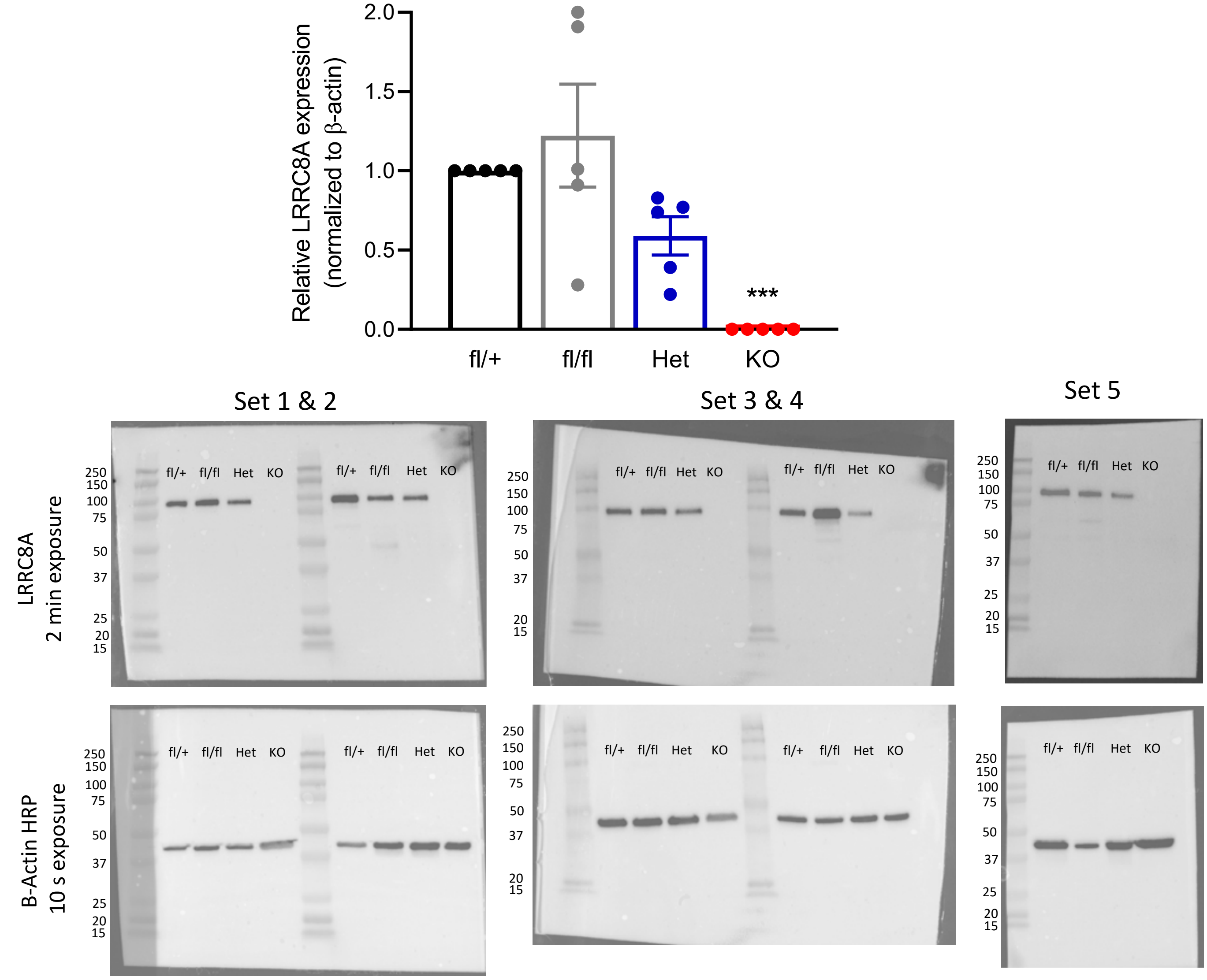


**Quantitative analysis and full-length western blot images for LRRC8A protein and β-actin probed in primary astrocyte culture lysates, prepared from fl/+, fl/fl, Het, or KO mice.** PAGE-separated proteins (10 µg/lane) were electrotransfered onto PDVF membrane, probed with monoclonal anti-LRRC8A antibody (Santa Cruz, cat #sc-517113, 1:500) and matching secondary antibody, and visualized using an ECL kit and BioRad Gel Doc Imager. The membranes were then stripped, re-probed with the HRP-conjugated anti-β-actin antibody (Sigma Millipore, cat #A-3854 , 1:50,000) and processed in a similar manner.

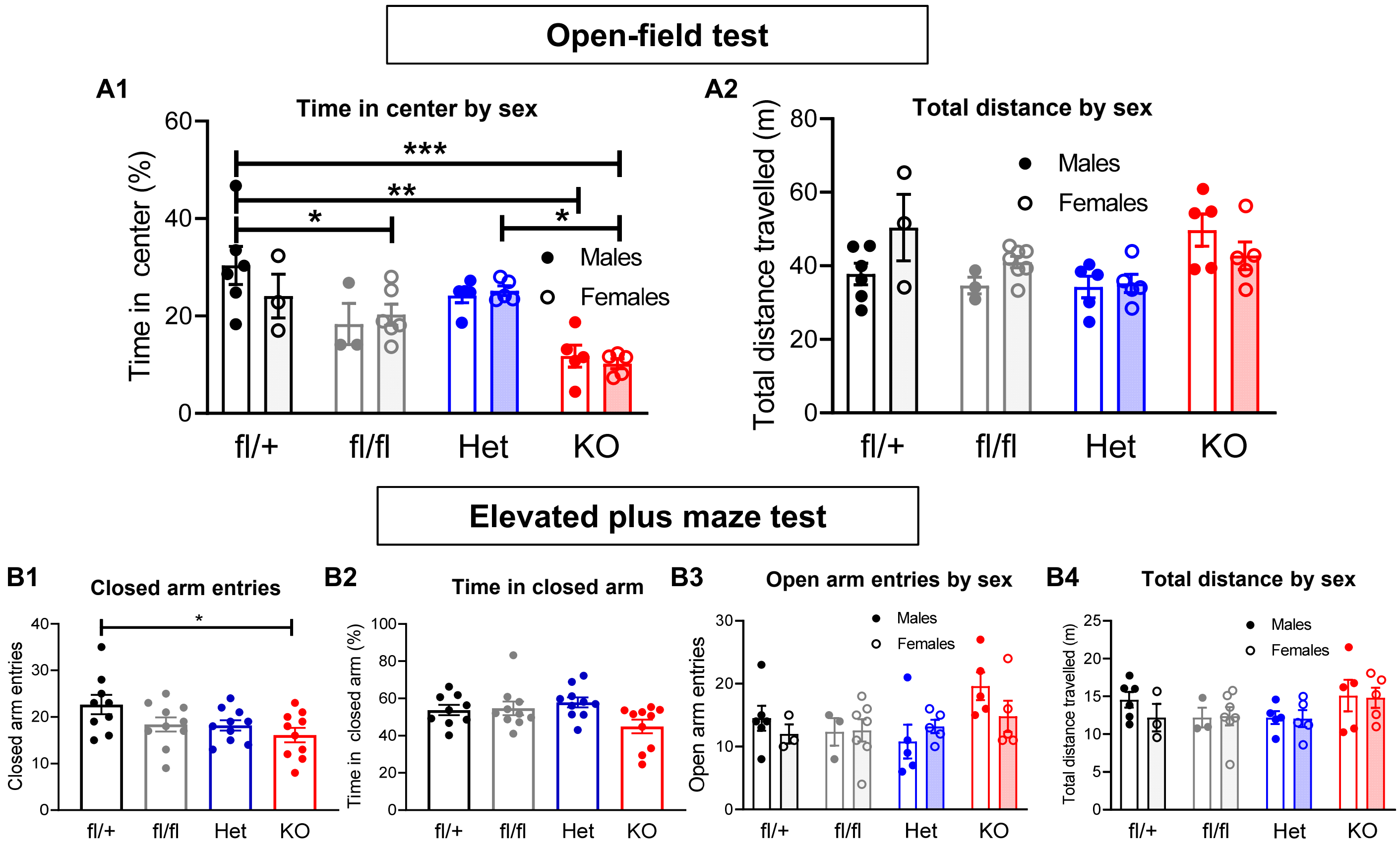


**Extended behavioral analysis of possible sexual dimorphism in behavioral tests performed in LRRC8A KO mice and their littermate controls.** (A) The Open Field Test (OFT) in 6-week-old male [solid symbols] and female [open symbols] mice. The observations were done over a 10-min period in 9 fl/+ and 10 mice in all other genotypes. The time in the center (A1) and the total distance traveled (A2), as compared by sex in fl/+ (6 male, 3 female), fl/fl (3 male, 7 female), Het (5 male, 5 female), and KO (5 male, 5 female) mice. Analyzed by two-way ANOVA with Bonferroni correction. (B) The Elevated Plus Maze (EPM) test performed in the same groups. Animal behavior was recorded for 5 min. The number of closed arm entries (B1) and the percentage of time spent in the closed arms (B2) are shown combined for both sexes. These data represent additional analyses for the experiments shown in Fig. 3 of the main manuscript. The number of open arm entries (B3) and total distance travelled (B4) was further analyzed by sex in fl/+ (6 male, 3 female), fl/fl (3 male, 7 female), Het (5 male, 5 female), and KO (5 male, 5 female) mice. In B3 and B4 solid symbols represent males and open symbols correspond to females. Data are the mean values ±SEM of 9 fl/+ mice and 10 mice in other groups, analyzed by one-way ANOVA (B1, B2) or two-way ANOVA (B3, B4) with Bonferroni correction. No significant differences were found between sexes. Combined data for both sexes for the closed arm entries were different between fl/+ and KO mice, $^*p<0.05$.

### C.S. Wilson *et al.* ● SUPPLEMENTARY FIGURE 7.1

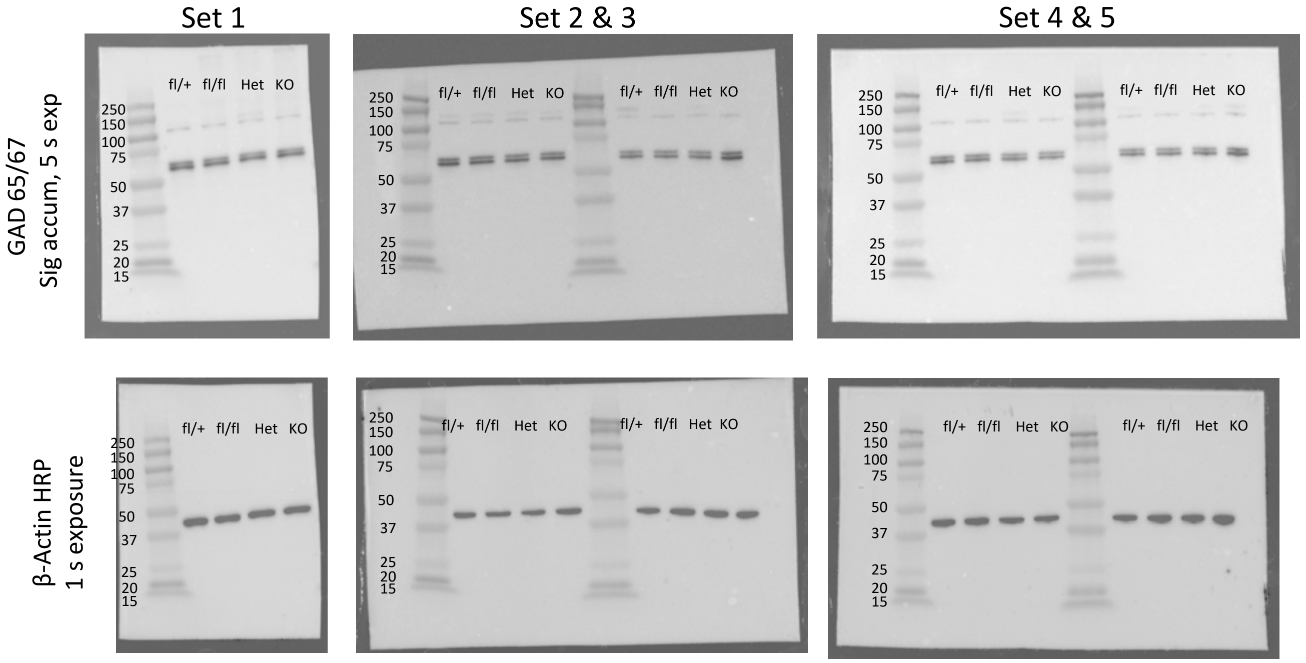


**Full-length western blot images for GAD65/67 protein and β-actin immunoreactivity probed in five sets of whole-brain tissue lysates from fl/+, fl/fl, Het, or KO mice.** PAGE-separated proteins (5 µg/lane) were electrotransfered onto PDVF membrane, probed with polyclonal anti-GAD65/67 (Millipore-Sigma, cat #G5163, 1:2,000), and matching secondary antibody and visualized using an ECL kit and BioRad Gel Doc Imager. The membranes were then stripped, re-probed with the HRP-conjugated anti-β-actin antibody (Sigma Millipore, cat #A-3854 , 1:50,000), and processed in a similar manner.

### C.S. Wilson *et al.* ● SUPPLEMENTARY FIGURE 7.2

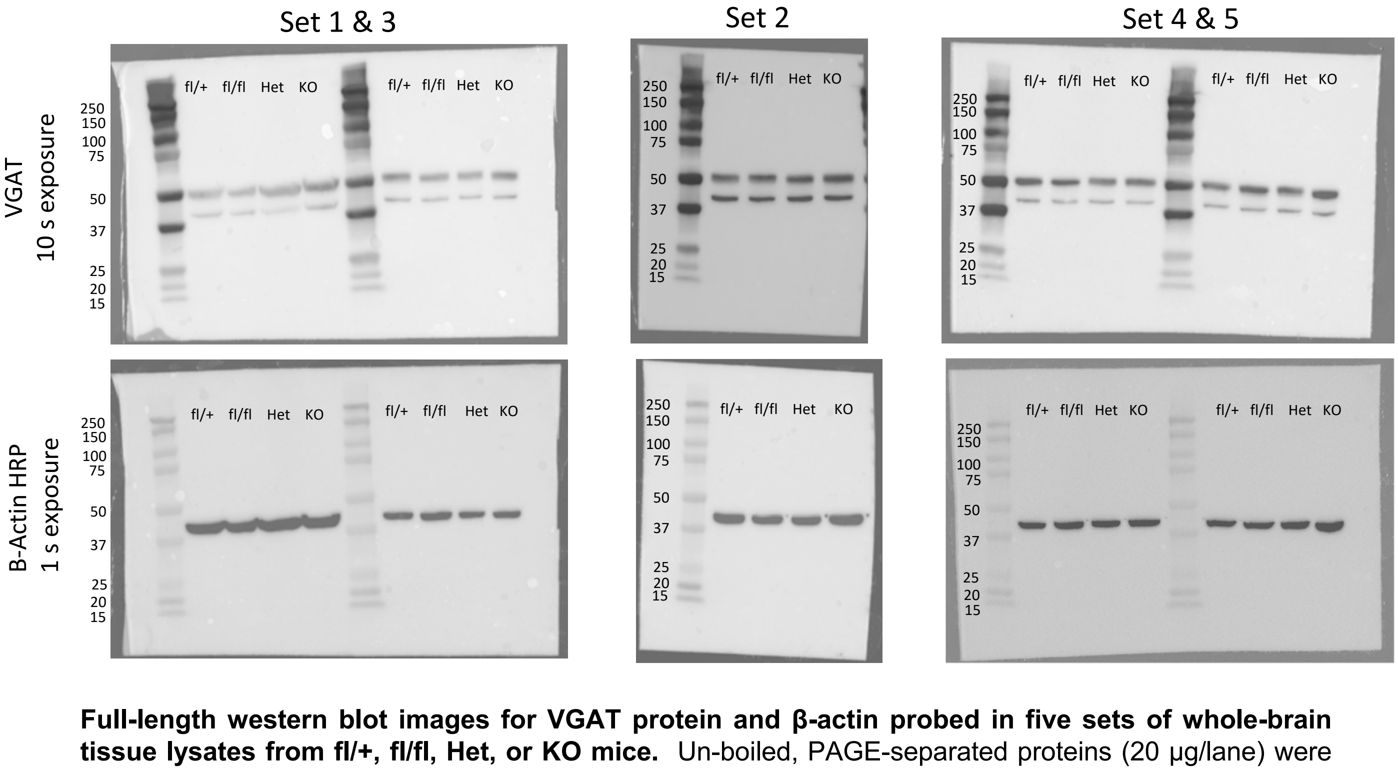


**Full-length western blot images for VGAT protein and β-actin probed in five sets of whole-brain tissue lysates from fl/+, fl/fl, Het, or KO mice.** Un-boiled, PAGE-separated proteins (20 µg/lane) were electrotransfered onto PDVF membrane, probed with polyclonal anti-VGAT (Invitrogen, cat. #Pa5-27569, 1:1,000), and matching secondary antibody and visualized using an ECL kit and BioRad Gel Doc Imager. The membranes were then stripped, re-probed with the HRP-conjugated anti-β-actin antibody (Sigma Millipore, cat #A-3854 , 1:50,000), and processed in a similar manner.

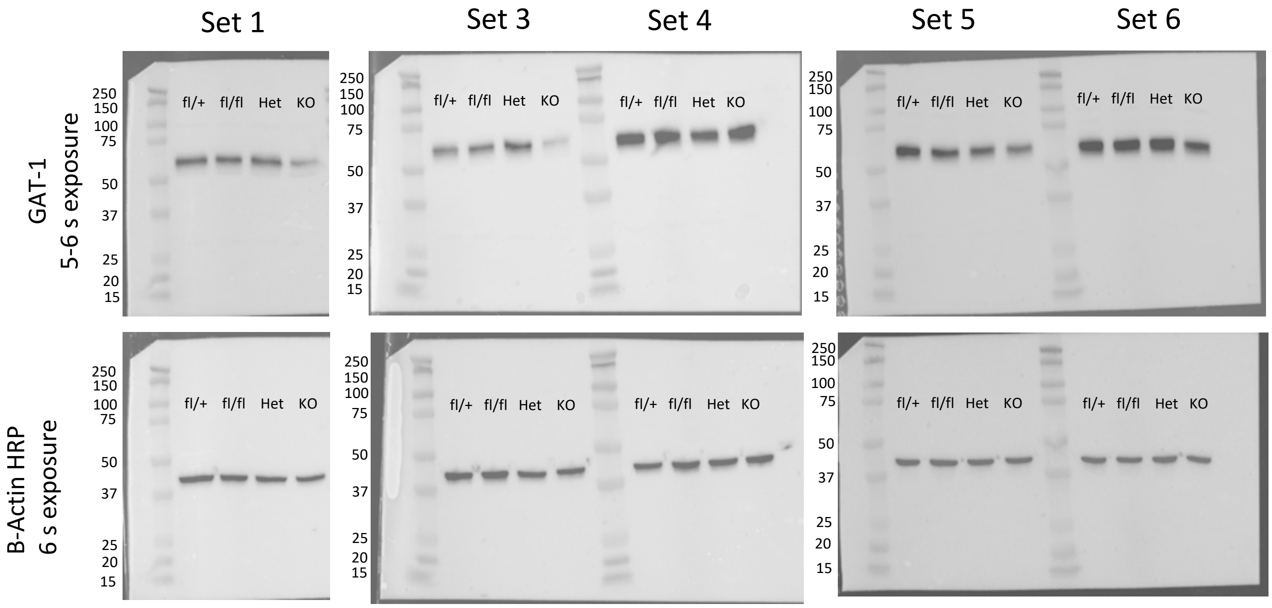


**Full-length western blot images for GAT-1 protein and β-actin probed in five sets of whole-brain tissue lysates from fl/+, fl/fl, Het, or KO mice.** Un-boiled, PAGE-separated proteins (10 µg/lane) were electrotransfered onto PDVF membrane, probed with polyclonal anti-GAT-1 (Abcam, cat. #ab72448, 1:500), and matching secondary antibody and visualized using an ECL kit and BioRad Gel Doc Imager. The membranes were then stripped, re-probed with the HRP-conjugated anti-β-actin antibody (Sigma Millipore, cat #A-3854 , 1:50,000), and processed in a similar manner.

C.S. Wilson *et al.* ● **SUPPLEMENTARY FIGURE 8.1**

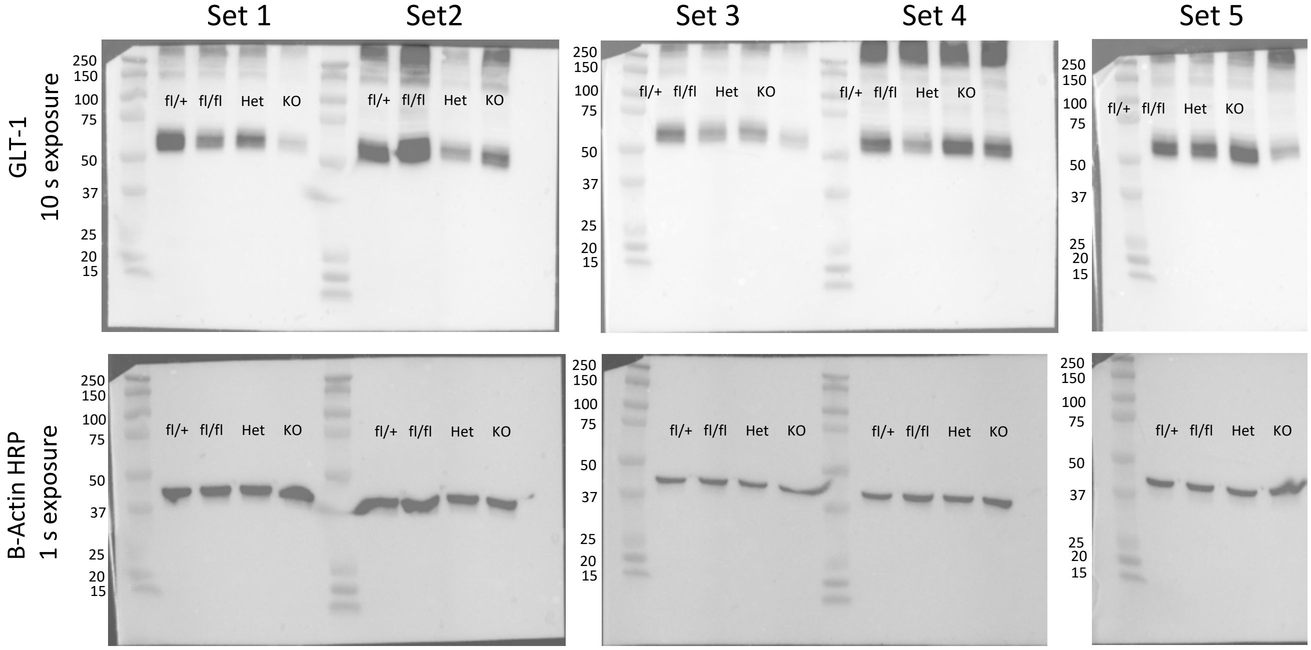


**Full-length western blot images for glutamate transporter GLT-1 and β-actin immunoreactivity probed in five sets of whole-brain tissue lysates from fl/+, fl/fl, Het, or KO mice.** PAGE-separated proteins (5 µg/lane) were electrotransfered onto PDVF membrane, probed with polyclonal anti-GLT-1 (Abcam, cat. #ab41621, 1:10,000) and matching secondary antibody and visualized using an ECL kit and BioRad Gel Doc Imager. The membranes were then stripped, re-probed with the HRP-conjugated anti-β-actin antibody (Sigma Millipore, cat. #A-3854 , 1:50,000), and processed in a similar manner.

C.S. Wilson *et al.* ● **SUPPLEMENTARY FIGURE 8.2**

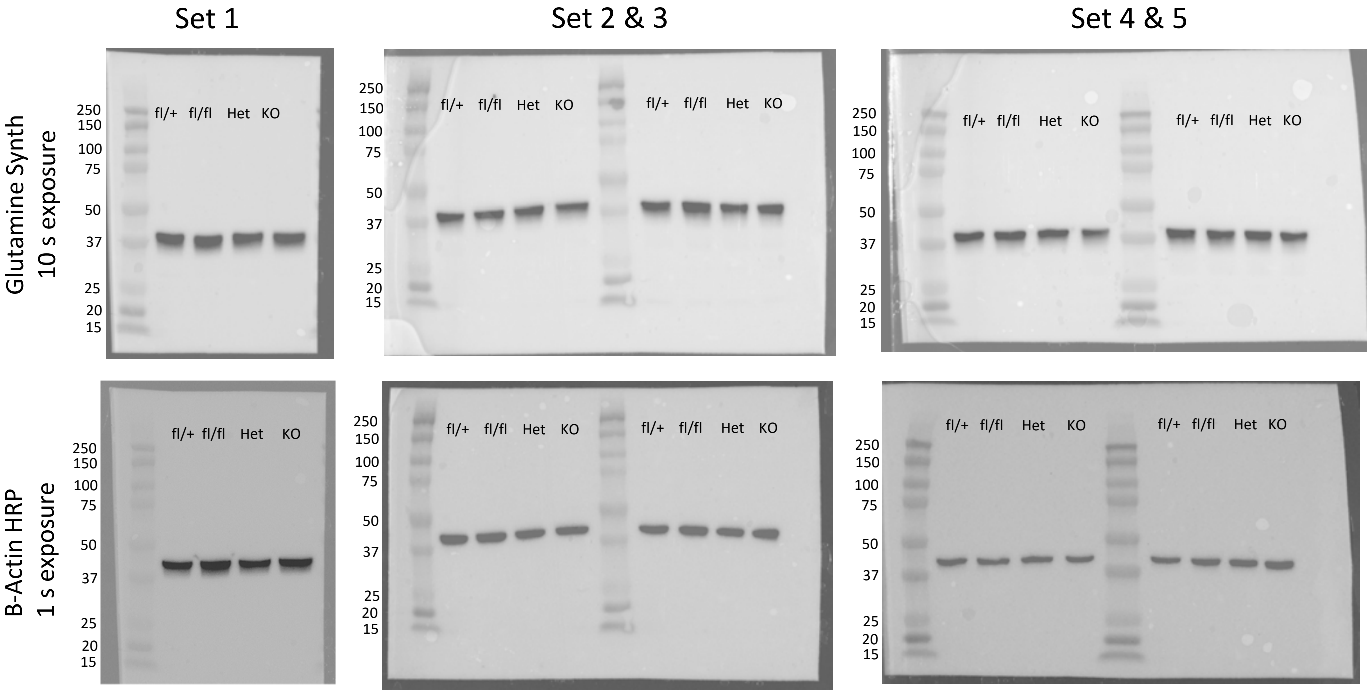


**Full-length western blot images for glutamine synthetase (GS) and β-actin immunoreactivity probed in five sets of whole-brain tissue lysates from fl/+, fl/fl, Het, or KO mice.** PAGE-separated proteins (10 µg/lane) were electrotransfered onto PDVF membrane, probed with polyclonal anti-GS (Millipore-Sigma, cat. #G2781, 1:20,000) and matching secondary antibody and visualized using an ECL kit and BioRad Gel Doc Imager. The membranes were then stripped, re-probed with the HRP-conjugated anti-β-actin antibody (Sigma Millipore, cat. #A-3854 , 1:50,000), and processed in a similar manner.
